# An integrated transcriptomic cell atlas of human endoderm-derived organoids

**DOI:** 10.1101/2023.11.20.567825

**Authors:** Quan Xu, Lennard Halle, Soroor Hediyeh-zadeh, Merel Kuijs, Umut Kilik, Qianhui Yu, Tristan Frum, Lukas Adam, Shrey Parikh, Manuel Gander, Raphael Kfuri-Rubens, Dominik Klein, Zhisong He, Jonas Simon Fleck, Koen Oost, Maurice Kahnwald, Silvia Barbiero, Olga Mitrofanova, Grzegorz Maciag, Kim B. Jensen, Matthias Lutolf, Prisca Liberali, Joep Beumer, Jason R. Spence, Barbara Treutlein, Fabian J. Theis, J. Gray Camp

## Abstract

Human stem cells can generate complex, multicellular epithelial tissues of endodermal origin *in vitro* that recapitulate aspects of developing and adult human physiology. These tissues, also called organoids, can be derived from pluripotent stem cells or tissue-resident fetal and adult stem cells. However, it has remained difficult to understand the precision and accuracy of organoid cell states through comparison with primary counterparts, and to comprehensively assess the similarity and differences between organoid protocols. Advances in computational single-cell biology now allow the integration of datasets with high technical variability. Here, we integrate single-cell transcriptomes from 218 samples covering organoids of diverse endoderm-derived tissues including lung, pancreas, intestine, liver, biliary system, stomach, and prostate to establish an initial version of a human endoderm organoid cell atlas (HEOCA). The integration includes nearly one million cells across diverse conditions, data sources and protocols. We align and compare cell types and states between organoid models, and harmonize cell type annotations by mapping the atlas to primary tissue counterparts. To demonstrate utility of the atlas, we focus on intestine and lung, and clarify ontogenic cell states that can be modeled *in vitro*. We further provide examples of mapping novel data from new organoid protocols to expand the atlas, and showcase how integrating organoid models of disease into the HEOCA identifies altered cell proportions and states between healthy and disease conditions. The atlas makes diverse datasets centrally available, and will be valuable to assess organoid fidelity, characterize perturbed and diseased states, and streamline protocol development.

## Introduction

It is a major goal to model complex aspects of human tissues in controlled conditions. Such in vitro human model systems can be used as inroads into human-specific biology and disease, as well as accurate alternatives to animal models (Loewa et al., 2023). Organoids are 3D cell cultures derived from pluripotent, fetal, or adult stem cells (PSCs, FSCs, ASCs) that recapitulate important aspects of cell composition, cytoarchitecture, and functional properties of the tissue counterpart (Kim et al., 2020). Organoid protocol development and engineering methodologies are rapidly emerging, such that there is strong variation in culture conditions and stem cell origin. It remains challenging to assess how well the complexity of cell states and interactions that occur in vitro reflect counterparts in vivo. The rapidity of organoid protocol development makes it difficult to compare between studies and protocols, hampering the assessment of correlation between previously generated and newly presented organoid datasets. An additional challenge is that datasets are not centrally compiled and protocols are not consistently reported, which makes it difficult to assess the similarity of different organoids to primary tissue, identify off-target or missing cell types, and predict genetic drivers that guide cell differentiation (Camp et al., 2018). Overcoming these obstacles would lead to new insight into basic human biology, as well as new opportunities for therapy development and rigor in clinical studies (Rozenblatt-Rosen et al., 2017; Bock et al., 2020). Advances in technology have led to the growth of single-cell transcriptome datasets, both in terms of dataset size and quantity. This has prompted collaborations to create extensive reference atlases for adult and developing human organs (Rozenblatt-Rosen et al., 2017; Bock et al., 2020; Tabula Sapiens Consortium* et al., 2022; Jain et al., 2023; Han et al., 2020). Organoids offer the opportunity to deepen our understanding of health and disease, by providing avatars of diverse developmental stages, genetic variation, and disease states that will complement primary tissue atlases (Bock et al., 2020). However, the scale of generating a comprehensive organoid atlas within individual research groups is currently impractical. Therefore, the integration of datasets generated by the wider research community becomes crucial.

The endoderm represents the innermost of the three primary embryonic germ layers formed during gastrulation. It contributes to the development of the epithelial lining of a variety of different organs including thyroid, esophagus, lung, pancreas, liver, biliary system, stomach, small intestine and colon (Zorn and Wells, 2009). Complex endodermal three-dimensional (3D) organoids from epithelial ASCs were first established after the discovery of tissue-resident intestinal stem cells (ISCs) marked by the expression of Lgr5 (Barker et al., 2007). In vivo, these stem cells reside in the intestinal crypt niche and underlie the renewal of the intestinal epithelium. Lgr5+ ASCs can be isolated and propagated in vitro in a 3D matrix to form organoids where media composition such as growth factors can promote stem cell proliferation and differentiation into intestinal epithelial cell types (Fujii et al., 2018). Tissue-resident ASCs have been discovered in a variety of endoderm-derived adult epithelial tissues, and protocols with specific combinations and concentrations of growth factors have been developed to propagate and differentiate ASCs into diverse cell types of corresponding tissues (Kim et al., 2020). Protocols have also been developed to cultivate FSCs from developing tissues (Sprangers et al., 2021) or to differentiate PSCs into various endodermal tissue types, enabling exploration of human ontogenetic processes (McCauley and Wells, 2017). In order to calibrate organoid fidelity, it is necessary to have a relative comparison of the cell state landscapes across the diversity of organoid protocols and sources.

Here, we present an integrated single-cell transcriptomic atlas of human endoderm-derived organoids encompassing nine different tissues, combining data from 55 publications and 218 individual single-cell sequencing experiments (datasets), comprising a total of 806,646 cells from ASC, FSC, iPSC, and ESC-derived organoids. After benchmarking multiple integration approaches, we integrated the datasets using a label-aware deep representation learning approach, and then performed re-annotation to establish a consensus cell type reference, enhancing our understanding of the endoderm-derived organoid systems and their relevance for disease modeling. Using intestine and lung organoids as examples, we compared cell states from diverse stem cell origins, uncovering insights into the functional relevance of these organoid systems. We applied the atlas to integrate and assess multiple new organoid protocols (129,254 cells from 2 tissues, 5 datasets, and 31 samples), validating the atlas as a valuable resource for analyzing future organoid data and expanding our knowledge of organoid development. We use the atlas to map data from organoid models of disease, which facilitated rapid assessment and comparison of control and diseased samples. Overall, our study provides a roadmap for constructing and utilizing organ and organoid cell atlases to explore human-specific biology.

## Results

### Data integration constructs the organoid atlas for endoderm-derived organs

To create an endoderm-derived organoid cell atlas, we assembled single-cell RNA sequencing (scRNA-seq) and single-nucleus RNA-seq (snRNA-seq) data from 55 datasets, including 1 newly generated dataset containing 45,281 cells collected from 11 separate samples (small and large intestine, stomach and liver organoids) and 54 published datasets (Fig. 1a, Extended Data Table 1). These datasets encompass samples from 218 experiments conducted on organoid models of nine different organs (lung, liver, biliary system, stomach, pancreas, small and large intestine, prostate, salivary glands) (Fig. 1b). Data were obtained using multiple scRNA-seq protocols, including plate-based methods such as Smart-seq, CEL-seq, and Sort-seq, as well as commercialized droplet-based methods (e.g. 10X Genomics) (Fig. 1c). Based on availability, we incorporated organoid datasets that model healthy states primarily of human endoderm-derived tissues, with source material from ESCs, iPSCs, FSCs, or ASCs (Fig. 1d). Notably, we obtained data of each stem cell source from intestine, lung, liver, and biliary system organoid models (Fig. 1b and d). In total, we collected 806,646 cells to be utilized for downstream integration and analysis (Fig. 1a-d). This extensive cell collection will serve as a valuable resource, providing opportunities to explore diverse technical and biological phenomena and extract meaningful insights through comparisons across datasets.

**Figure 1.**
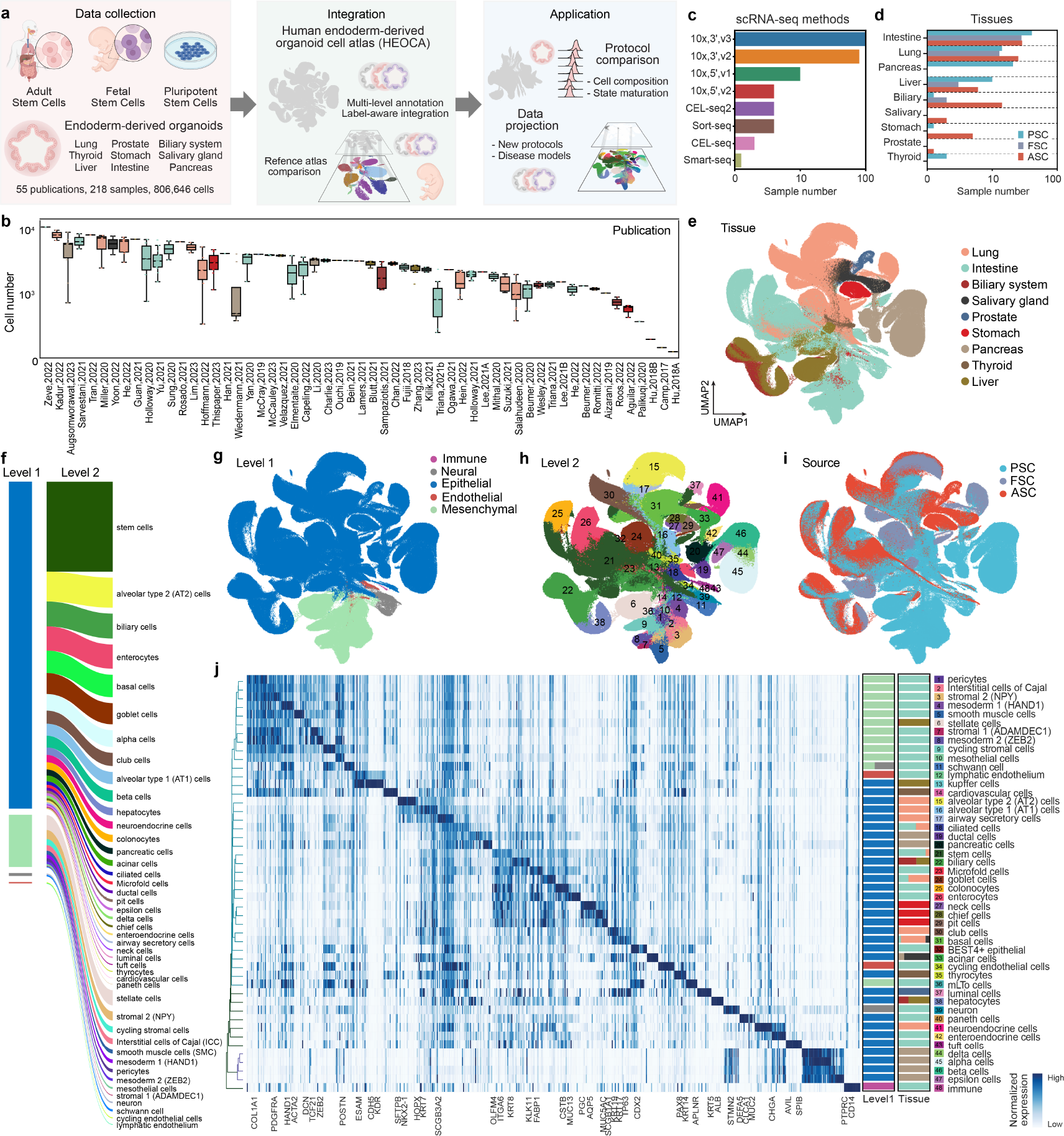
Integrated transcriptome cell atlas of human endoderm-derived organoids. (a) Schematic overview of the atlas integration and downstream analyses. (b) Boxplot showing the number of cells of each sample in all publications. (c-d) Barplot showing the number of samples grouped by different single cell sequencing methods (c) and by tissue and stem cell source organoid (d), respectively. (e) UMAP of the organoid atlas colored by tissue. (f) Overview of level 1 (colormap see (g)) and level 2 cell annotations and cell proportion. (g-i) Organoid atlas by level 1 annotations (g), level 2 annotations (h) or by stem cell source (i). (j) Heatmap showing marker gene expression for each level 2 cell type in the atlas. Side stacked barplots show proportions of cell types at level 1 and tissue type annotations.

Consensus cell type definitions from scRNA-seq data across studies are currently insufficient, particularly for different organoid developmental states. To define organoid cell types, we employed an annotation method that first clusters cells in each dataset at a high resolution and assigns cluster annotations based on known marker gene expression and differential expression compared to other clusters (Extended Data Table 2). To assist with label-aware integration, we established a three-level hierarchical cell type annotation (level 1 = class, level 2 = type, level 3 = subtype) (Extended Data Fig. 1). To effectively address batch effects and achieve a robust integration of the atlas, we assessed nine different data integration methods using single-cell integration benchmarking on ten randomly chosen samples (Polański et al., 2020; Xu et al., 2021; Lopez et al., 2018; Korsunsky et al., 2019; He et al., 2020; Büttner et al., 2019; De Donno et al., 2023) (Extended Data Fig. 1a). Aggregate scores based on batch correction and conservation of biological structure suggested that the label-aware integration method scPoli (Deprez et al., 2020; Luecken et al., 2022) achieved the most successful integration (Extended Data Fig. 1a-b). Utilizing scPoli, we generated an integrated embedding that encompassed all organoid cells, enabling a cohesive representation of the diverse data. This integration overcame batch effects (batch correction = 0.43 and biology conservation = 0.98) and generated a unified view of the HEOCA (Fig. 1e, Extended Data Fig. 1b). We reannotated high-resolution clusters of the integrated atlas based on the most frequently occurring cell type per cluster resulting in 5 cell classes at level 1, 48 cell types at level 2, and 51 cell subtypes at level 3 (Fig. 1f-h, Extended Data Fig. 1c-d).

Overall, we observed that epithelial cells from different organs clustered together within the integrated atlas, and that many clusters were composed of cells from different stem cell sources (Fig. 1e and i). This observation suggests different cell types in the same organ model from different protocols share more similarity compared to cell types from different types of organoids. However, we also identified certain cell types with contribution from multiple organoid models. For example, goblet cells were found in both intestine (67.05%) and lung (32.95%), with a minor presence in other organs (0.01%). Similarly, basal cells were observed in the lung (76.59%), salivary gland (12.52%), intestine (9.11%), and thyroid (1.25%) models (Fig. 1j). These results suggest the existence of cell types that exhibit partial or shared characteristics across different organ models within the atlas, and also may indicate off-target cells within organoids. We identified markers for each integrated cell type, which could be used to distinguish cell types across the diverse datasets and protocols (Fig. 1j, Extended Data Table 3). For example, OLFM4 and TP63 marked stem cells and basal cells, respectively, across organoid models (Fig. 1j). We note that there are cases that were difficult to explain, where cells derived from organoid models of a certain organ clustered with cells annotated as from a different organ. Given the difficulty in precisely controlling organoid development, especially PSC-derived organoids, it is known that there can be off-target cells in organoids (Yu et al., 2021). For instance, basal cells are not observed in adult human intestinal tissues. However, our analysis revealed that approximately 9% of basal cells within HEOCA originate from intestinal organoids (Fig. 1j). This observation suggests that organoids can contain cell types that are not the desired target for a specific organoid protocol (off-target cells) alternatively, there could also be instances where organoids derived from primary FSCs or ASCs are contaminated due to handling or to adjacency of tissues during the tissue acquisition, or cell states could be different from the tissue of origin due to stem cell plasticity. Therefore, it is important to develop strategies to compare organoid cells to reference counterparts.

### Reference atlas comparison to assess organoid fidelity

To evaluate cell states observed in the HEOCA, we obtained published scRNA-seq data on human endoderm-derived organs from adult (small and large intestine, lung, liver, pancreas, prostate, salivary gland) (Tabula Sapiens Consortium* et al., 2022) (Fig. 2a-b) and fetal (small and large intestine, lung, liver, pancreas, stomach) (Cao et al., 2020) (Fig. 2c-d) specimens. In adult and fetal datasets, we identified major cell types from each tissue (Fig. 2a-d), and then compared organoid cell types and states to these primary tissue references using neighborhood graph correlation (Ton et al., 2023). This analysis revealed a spectrum of maximum correlation values to neighborhoods found in the adult and fetal datasets, with a general higher similarity to adult tissue cell types (Fig. 2e, Extended Data Fig. 2). We quantified the proportion of cell types in each organoid sample and compared the similarity of each cell type to counterparts in the adult and fetal tissue (Fig. 2f-h) and found that ASC-derived and PSC-derived organoids had the highest similarity to adult and fetal counterparts, respectively, with FSC-derived organoid cell states having an intermediate distribution (Fig. 2i).

**Figure 2.**
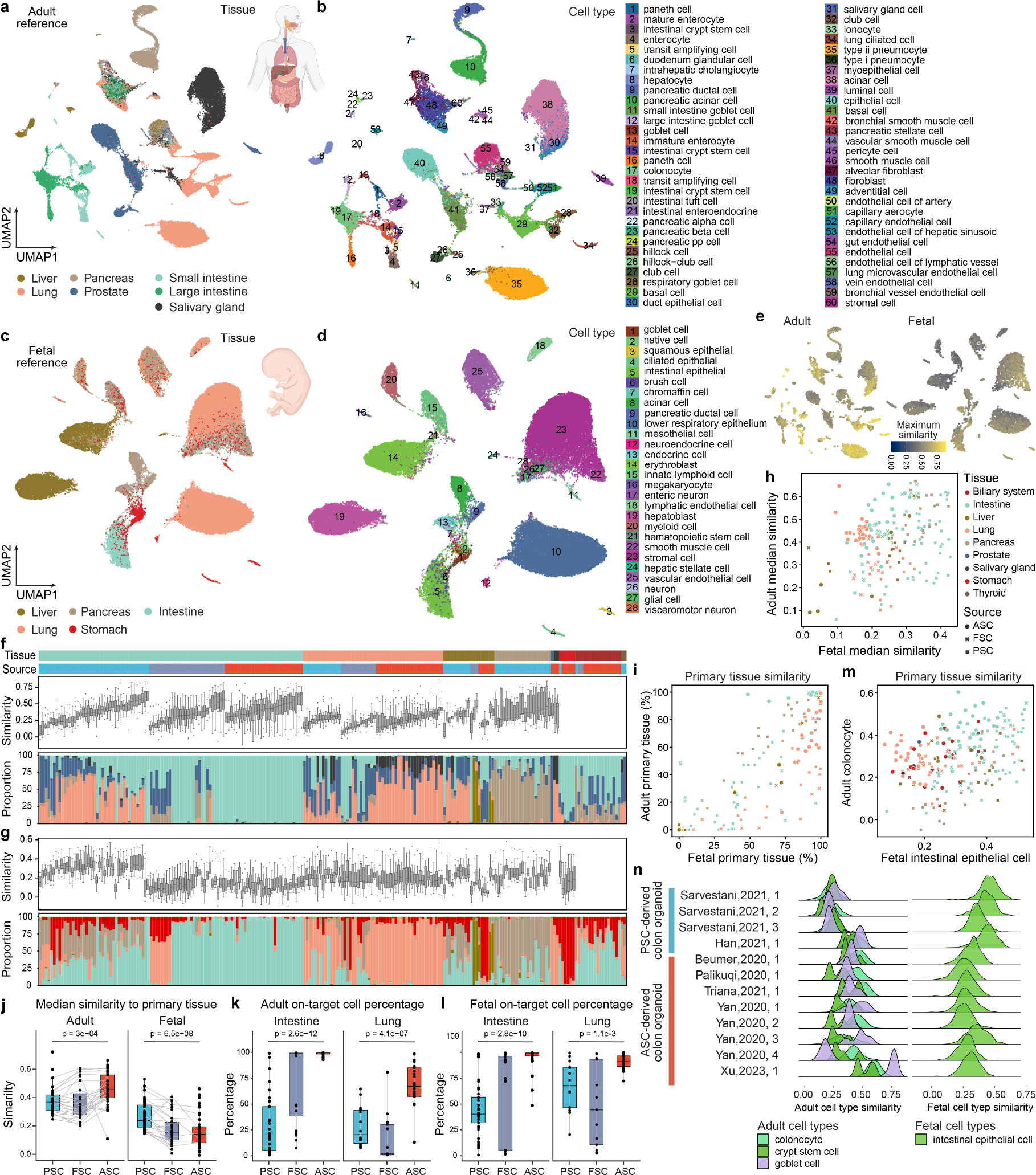
Mapping organoid cell types to a primary tissue reference atlas to assess organoid fidelity. (a-b) UMAP representations are shown for an integrated object comprising primary adult tissues (a) and cell types (b), as presented in the original publication. (c-d) Similarly, UMAP representations are displayed for an integrated object containing primary fetal tissues (c) and cell types (d) from the original publication. (e) UMAPs for primary adult (left) and fetal (right) tissues demonstrate the maximum similarity of all organoids within the comprehensive cross-tissue organoid atlas. (f) A boxplot illustrates the similarity of all cell types within corresponding primary adult tissues. The accompanying barplot indicates the tissue proportion of the most similar adult tissue. Samples are sorted by organoid tissue and the source of stem cells. The upper annotations indicate the organoid tissue and the source of stem cells. (g) The similarity of all cell types within corresponding primary fetal tissues is depicted, with a corresponding barplot showing the tissue proportion of the most similar fetal tissue. Samples are sorted as in (f). (h-i) Scatter plots display the median similarity of all cell types (h) and on-target tissue percentages (i) within corresponding primary adult and fetal tissues. (j) Boxplots present the median similarity to primary adult (left) and fetal (right) cell types among different sources of stem cell organoids. Gray lines connect the same cell types from different stem cell sources. (k) Boxplots reveal the adult on-target cell percentage in the intestine (left) and lung (right) organoid samples, ordered as ASC, FSC, and PSC from left to right, respectively. (l) The same analysis as (k) on fetal tissues. (m) Tissue similarity is shown for all organoid samples compared to adult colonocytes and fetal intestinal epithelial cells. (n) Ridge plots display all colon samples similar to colon primary tissue cell types, with adult and fetal cell types arranged from left to right.

To assess off-target cells within organoids, we projected organoid cells to the fetal and adult primary tissue atlases, and calculated the majority tissue type of the 100 nearest primary tissue neighbors, normalized by total number of cells of a given tissue type. Our analysis revealed that PSC-derived organoids have a lower on-target percentage in both fetal and adult primary tissues compared with FSC- and ASC-derived organoids (Fig. 2g-h, and j). For more specific downstream analyses, we focused on intestine and lung organoids, where there are multiple datasets from different stem cell sources. FSC and ASC-derived intestine organoids demonstrated high on-target percentages, with an average of 80% in FSC-derived organoids and 99% in ASC-derived organoids (Fig. 2k). In contrast, PSC-derived organoids displayed median on-target percentage of between 25% and 50% depending on fetal or adult reference atlas comparison, however this is likely a low estimate as datasets from early organoid time points are difficult to assess using this reference comparison metric (Fig. 2k). In the lung organoids, we observed a median on-target percentage of 75% in ASC-derived organoids. In contrast, PSC and FSC-derived organoids exhibited lower on-target percentages (Fig. 2k). Notably, the on-target percentage of intestine organoids in relation to fetal primary tissue closely mirrored that of adult tissue, with percentages of 55%, 80%, and 99% in PSC, FSC, and ASC-derived organoids, respectively (Fig. 2l). Focusing on colonic epithelial cell types, ASC-derived organoids exhibited a higher similarity with adult colonocytes, while PSC-derived intestine organoids showed a higher similarity with fetal intestinal epithelial cells, and we note that there is variation in maturation levels between protocols (Fig. 2m-n).

### Integrated intestinal organoid atlas reveals developing and adult physiology

To explore organoid cell states of different stem cell origin, we focused on intestinal organoid models, where there is substantial coverage from PSC, FSC, and ASC-derived organoid cells. This subset consisted of 98 samples from 23 different publications representing 353,140 single cell transcriptomes of duodenum, ileum, and colon tissues (Fig. 3a, Extended Data Fig. 3a, Extended Data Table 1). We re-integrated all cells and defined five cell types at level 1, 26 cell types at level 2, and 32 cell types at level 3 within the atlas (Fig. 3b, Extended Data Fig. 3b-c). This integrated intestinal organoid atlas (HIOCA) covers epithelial states from the duodenum, ileum, and colon and PSC-derived organoids additionally contain a large fraction of mesenchymal cells, and minor populations of neural, endothelial, and immune cell types (Fig. 3b-c). The integrated atlas provides opportunities to explore differentiation trajectories across protocols, and could be used to identify variant and invariant gene expression profiles. (Extended Data Fig. 3d-f).

**Figure 3.**
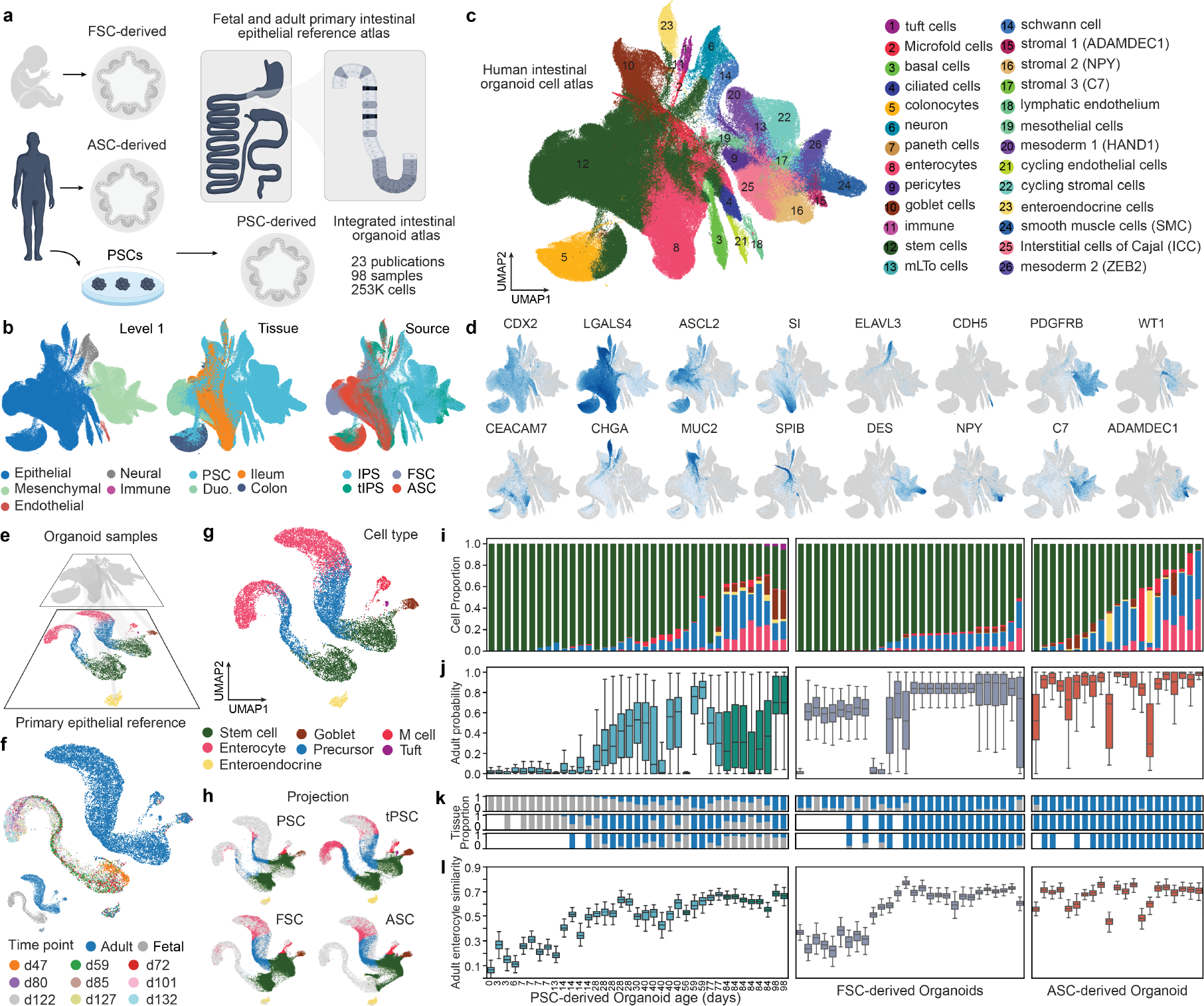
Human intestinal organoids from different stem cell origins generate developing and adult cell states. (a) Analytical design of the intestine organoid sub atlas and the comparison to the primary reference tissue. (b) UMAP of healthy intestinal organoid atlas colored by level 2 cell type annotation. (c) UMAP of healthy intestinal organoid atlas colored by level 1 cell type annotation, source of intestine tissues, and source of stem cells. (d) UMAP showing intestine cell type marker gene expression. (e) Analytical design of the intestine organoid sub atlas and the comparison to the primary reference tissue. (f-g) UMAP of the integrated intestine fetal and adult primary tissue single-cell object colored by adult sample or fetal sample age (f) and cell type (g). (h) Project the intestine organoid atlas cells onto fetal and adult primary epithelial single-cell objects categorized by PSC, tPSC(transplant PSC), FSC, and ASC-derived organoid samples. (i) Barplot illustrates the predicted cell proportions of each organoid sample mapped to the primary tissue objects. The samples are divided by PSC, FSC, and ASC-derived organoid samples, with PSC-derived organoids further ordered by organoid age and FSC/ASC-derived organoids ordered by the percentage of stem cells. (j) Boxplot shows the predicted probability of cell mapping to adult samples. (k) Barplots illustrate the predicted tissue (fetal: gray and adult: blue) proportions. From up to the bottom are stem cells, precursor enterocytes, and enterocytes. (l) Boxplot shows the adult enterocytes similarity of each organoid sample. (j) to (l) have the order of organoid samples consistent with (i).

In order to build a reference to assess intestinal organoid fidelity and maturation we next integrated time series scRNA-seq data of duodenal development (ten time points ranging from 59 days to 132 days after fertilization) with adult intestinal epithelium (Yu et al., 2021; Elmentaite et al., 2021)(Fig. 3e). The integrated reference atlas revealed distinct fetal and adult stem cell to enterocyte differentiation trajectories, while demonstrating similar cell states between fetal and adult in other epithelial cell types, such as goblet cells, tuft cells, M cells, and enteroendocrine cells (Fig. 3f-g). To validate the reliability of this integration, we downloaded an additional 18 fetal and 5 adult samples from two publications (Cao et al., 2020; Elmentaite et al., 2021) and mapped cells to the primary embedding (Extended Data Fig. 3g-j). The mapping indicated that an average of 95% of fetal stem cells and enterocytes aligned with the fetal trajectory, while an average of 98% of adult stem cells and enterocytes aligned with the adult trajectory (Extended Data Fig. 3g-j). This analysis confirms that the integration of primary adult and fetal intestine tissue scRNA-seq data can discern fetal and adult cell states, providing a resource for investigating organoid maturity. We used scPoli to query small intestine organoid epithelial cells to the primary reference data and observed distinct similarity patterns among different stem cell sources (Fig. 3h). Specifically, PSC-derived organoids exhibited a higher proportion of cells resembling fetal primary tissue, while FSC-derived and ASC-derived organoids displayed a high similarity to adult primary tissues (Fig. 3h). This finding supports previous observations that FSC-derived intestinal organoids may undergo a transition in their identity and lose fetal characteristics upon extended culture in vitro (Edgar et al., 2022). We provide information on the proportion of cell types (Fig. 3i), adult projection probability (Fig. 3j), fetal or adult projection proportion (Fig. 3k), and similarity to each adult and fetal cell type (Fig. 3l, Extended Data Fig. 3k) for each PSC, FSC, and ASC-derived organoid sample. Collectively, these metrics show substantial variation between intestinal organoid samples. For example, PSC-derived organoids increase in complexity and reference similarity over time in culture, and after xenografting into a mouse host for maturation, the organoids obtain higher cellular diversity and similarity to primary tissue differentiated enterocytes. In addition, in these datasets FSC-derived organoids presented less cell type diversity than ASC-derived organoids, and there was variation in diversity and similarity between datasets from the same stem cell source type. These findings reveal the diversity of cell composition and cell maturation within organoids from different sources, time points, and protocols.

### Lung organoid data uncovers differences between organoids of varying stem cell origins and shows specific correlation to primary tissue

To complement our analysis of intestinal cell states, we created and analyzed a singular integrated object of lung organoid cells, representing the second most abundant tissue type in the HEOCA. This subset atlas consisted of 225,490 cells obtained from 52 samples and 13 different publications (Fig. 4a). For this, we employed an approach similar to our intestinal organoid analysis, involving integration, re-clustering, and reannotation of these data (Fig. 4b). The integrated human lung organoid cell atlas (HLOCA) encompasses cell types and states derived from PSCs, FSCs, and ASCs (Fig. 4c). Predominantly composed of epithelial cells, the lung organoid samples are further divided into 9 distinct cell types through higher-resolution clustering (Extended Data Fig. 4i). The UMAP representation showed expected integration of the diverse publications and samples into a shared embedding (Ext. Data Fig. 4a-b) with undifferentiated stem cells positioned centrally surrounded by more differentiated cell types (Fig. 4b) exhibiting cluster-specific expression of canonical markers of lung epithelial cell types (Fig. 4e). Cell type composition analysis per organoid revealed cell type variation depending on the stem cell of origin, reflecting the diversity of culture and isolation protocols among the datasets in the atlas (Fig. 4d). Organoids derived from PSCs displayed a higher proportion of lowly-differentiated LGR5-, OLFM4and ASCL2-defined stem cells, which were largely absent in the ASC-derived organoids. In turn, organoids obtained via ASC-protocols frequently contained a relevant proportion of club cells, whereas a high incidence of goblet and neuroendocrine cells was primarily observed in samples produced using FSC-protocols (Fig. 4d). Overall, inter-publication and inter-sample differences in cell type composition suggest effects from protocol, media, and growth factors. We provide a detailed structured account of the publicly available information on the organoid datasets included in the atlas and have initiated a resource containing accumulated published data on protocol components and intervals, which can be linked to the shared HLOCA object.

**Figure 4.**
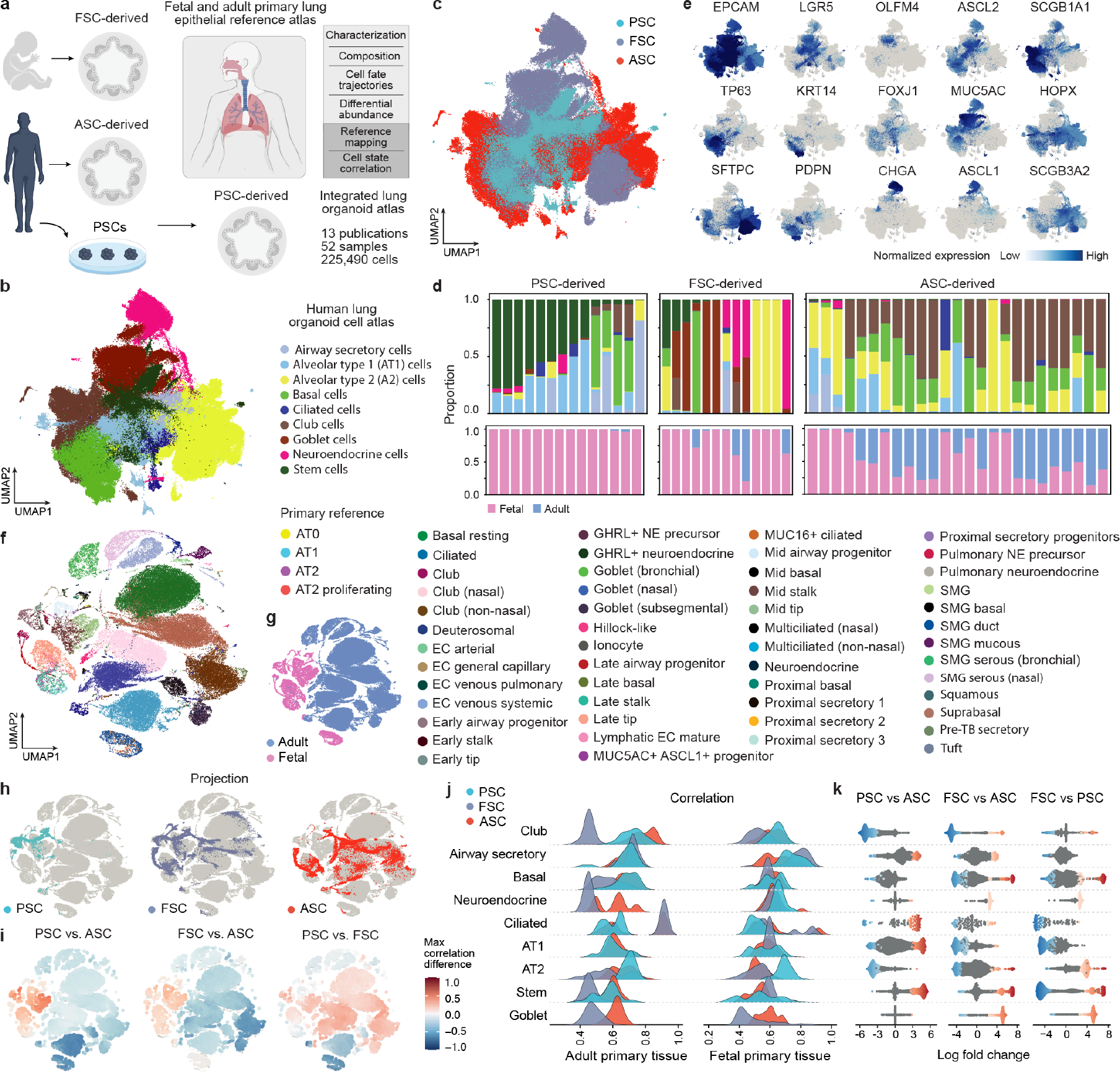
Human lung organoids from different stem cell origins generate developing and adult cell states. (a) Schematic of the analyses performed on the lung organoid sub atlas (HLOCA) and the comparison to the primary reference tissue. (b) UMAP of the integrated object of all lung organoid samples colored by cell type (c) UMAP of the integrated object of all lung organoid samples colored by stem cell of origin. (d) Upper row: Stacked barplots depicting cell type composition per sample by stem cell of origin (PSC/FSC/ASC), (ordered by amount of stem cells in sample from left to right, colored by cell type label from (c)); Lower row: predicted maturity in relation to the integrated lung primary adult and fetal reference embedding, (ordered by amount of stem cells in sample from left to right). (e) UMAPs depicting selected marker expression of cells in the HLOCA. (f) Shared lung primary adult and fetal reference embedding colored by original author annotation (g) Shared lung primary adult and fetal reference embedding colored by tissue maturity (h) Mapping results of HEOCA lung organoid datasets to a shared lung primary adult and fetal reference embedding; from above: maturity of reference, PSC-, FSCand ASC-mapping (i) Difference of indicated max correlation between neighborhoods of the HEOCA lung organoid subset and the shared primary and fetal lung reference embedding. (j) Ranked ridgeline plot, ordered by mean maximum correlation, showing max correlation distribution over neighborhoods between organoid atlas and fetal or adult reference per indicated cell state; ridges resolved by stem cell of origin. (k) Differentially abundant regions in organoids derived from different stem cells annotated by cell state; colored by significance.

Next, through trajectory analysis by diffusion pseudotime, we simulated developmental pathways from central stem cells towards predicted peripheral terminal cell states (Extended Data Fig. 5d-e). Within these trajectories, we identifed putative driver genes associated with selected terminal states for each cell type and charted their expression patterns across pseudotime (Extended Data Fig. 5f). Notably, the late-pseudotime expression peaks of this analysis aligned well with known canonical markers of terminal cell states, such as SFTPB and LPCAT1 in AT2s, LYZ in goblet cells, KRT15 in basal cells and LCN2 in club cells. Additional genes that were uncovered by this analysis, such as INSM1 and RAB26 in neuroendocrine cells, warrant further investigation into their roles in development. We conducted analogous analyses for the intestinal cells of the HIOCA and found several highly cell fate correlated genes previously reported to have an association with the developmental trajectory of the respective cell type (Extended Data Fig. 5a-c). This included, among others, genes coding for fatty acid and triglyceride binding proteins such as FAB1, FAB2 and RBP2 in enterocytes. For intestinal goblet cells, canonical markers like MUC2, SPINK4, CLCA1, and FCGBP were observed. In enteroendocrine cells, secretogranins (SCG2, SCG3) and transcription factors (NEUROD1, FEV) were present. Specific markers such as advillin were detected in Tuft cells (AVIL) and the ETS transcription factor SPIB was found in microfold cells.

To gain insights into how the assembled lung datasets correspond with primary tissue, we integrated a unified reference of primary adult and fetal tissues (He et al., 2022; Deprez et al., 2020) (Fig. 4f-g, Extended Data Fig. 4d). The query to reference mapping of our lung organoid data led to preferential integration of the PSC-derived and most of the FSC-derived organoids into the fetal portion of the reference (Fig. 4d, 4h). In turn, with the caveat of mapping uncertainty and distance to the reference being relatively high in some of the included samples (Extended Data Fig. 4g-h), a relevant proportion of the ASC-derived organoid cells showed an additional similarity to the adult cells present in the reference (Fig. 4d, 4h). We evaluated the correlation of differential neighborhoods derived from Milo (Dann et al., 2022) between HEOCA lung organoids and the adult and fetal reference (Extended Data Fig. 4e-f). Projecting the difference in correlation between PSCand ASCand between FSCand ASC-derived organoids on the neighborhood representation confirmed that ASC-derived organoids had the highest similarity to adult tissue (Fig. 4i). Conversely, PSC-derived and FSC-derived cells had a higher similarity to fetal cells, with PSC-derived cells possessing a slightly higher correlation to many regions of the combined reference embedding. To quantify these observations, we generated ranked ridgeline plots depicting cell type and organoid type specific correlations within the HEOCA lung subset in relation to adult and fetal reference neighborhoods (Fig. 4j) as well as correlations resolved by primary cell type and organoid type (Extended Data Fig. 4c). Our analysis indicated that, with the exception of ciliated and airway secretory cells, either ASCor PSC-derived organoids tended to resemble the primary reference cells more closely than FSC-derived ones across most lung organoid cell types, in relation to both adult and fetal tissue. While, in relation to adult tissue, club cells demonstrated the highest average correlation across all organoids, the peak of correlation was formed by a portion of the ASC- and FSC-derived ciliated cells. In examining the best-represented primary tissue cell types, our results reaffirmed the higher resemblance of fetal primary tissue cell types to PSC- and FSC- and of adult primary tissue cell types to ASC-derived organoids (Extended Data Fig. 4c). The highest scoring primary cell type, when compared against the different organoid types, were adult ciliated and different secretory cells for ASC-derived organoids, fetal ciliated and secretory cells for FSC-derived organoids and mid tip, mid stalk and early tip cells for PSC-derived organoids (Extended Data Fig. 4c).

Lastly, differential abundance analysis of the annotated cell types between the organoids detailed that PSC-derived organoid samples lack club, basal and AT2 cells compared to those derived from ASC protocols (Fig. 4k). While neuroendocrine and goblet cells were particularly found in FSC-derived organoid samples, the rest of the cell types were missing in samples derived from this stem cell compared to those from ASC protocols. Yet, FSC-derived organoid samples were found to have more AT2, club and basal cell states compared to PSC-derived samples, which are predominantly missing in samples generated by PSC differentiation protocols. In summary, these results and analyses offer a comprehensive integrated atlas for lung organoids (HLOCA) serving as a complement to the HEOCA for the study of lung 3D-cultures at a single tissue level. Moreover, by further downstream analysis of the gathered data, we identify differences in cell type composition and abundance, trajectories and associated genes, explore resemblances to fetal and adult primary tissue and provide novel information about the current representation of primary tissue cell types in lung organoids derived from various stem cell sources.

### Assessment of new protocols and extension of the organoid cell atlas with new datasets

New organoid protocols are emerging frequently and the HEOCA offers a unique opportunity to assess these organoids via single-cell transcriptomics. To incorporate new datasets into the organoid atlas, we developed an integration model based on scPoli that provides a harmonized annotation and toolkit for downstream analysis and exploration (Fig. 5a). To examine the performance of our model, we accessed an unpublished dataset from a scRNA-seq (10X Genomics) time course of colon organoid development (days 1, 3, 4, 5, 6, 7, 8, and 10), where organoids were grown in a new media combination (Fig. 5b) Oost et al. in revision. Cells were projected onto the HEOCA, and annotated based on label transfer of the nearest organoid neighbors (Fig. 5c). Analysis of predicted cell type proportions revealed a decrease in the proportion of stem cells from day 3 (99.69%) to day 10 (62.39%) concomitant with an increase in differentiated goblet cells (0.26% to 3.44%), enteroendocrine cells (0.04% to 7.43%), and colonocytes (0.02% to 33.41%) (Fig. 5d). It is noteworthy that the day 1 sample was in a recovery, regenerative stage after organoid passaging, and exhibited a higher proportion of mature cell types (Fig. 5d). Comparison of each cell to the adult and fetal reference atlas confirmed a temporal increase in adult cell type similarity (Fig. 5e-f), and integration analysis indicated near perfect cell type prediction as intestine-derived (Fig. 5g). We incorporated five additional datasets (three published (Frum et al., 2023; Conchola et al., 2023; Mitrofanova et al., 2023) and one unpublished), encompassing two additional colon ASC-derived organoid (colon 2 and colon 3) and two lung ASC-derived organoid protocols (lung 1 and lung 2) (Fig. 5h, Extended Data Fig. 6). In the colon 2 dataset, we examined colon organoids derived from three healthy individuals cultured with or without IL-22 for three days before analysis Maciag et al. in revision. HEOCA comparison revealed that control samples exhibited a higher proportion of mature colonocytes (5.72%) compared to the IL22 treated samples (1.53%) suggesting that the presence of IL-22 affects colonocyte differentiation (Fig. 5h-i, Extended Data Fig. 6a-c). In the colon 3 dataset, colonic epithelial tissue was generated through seeding human colon ASC-derived organoids on a scaffolded hydrogel within a fluidic chip (Mitrofanova et al., 2023). This dataset also provides two control samples culturing organoids in either matrigel or matrigel supplemented with 20% collagen. The integration analysis demonstrated that this new protocol led to the maturation of colonocytes, as indicated by a significantly higher proportion of colonocytes compared to the control samples (Fig. 5h-i, Extended Data Fig. 6d-f). Specifically, the day 4 control samples displayed a 30.32% proportion of colonocytes, compared with 48.20% and 66.14% at day 14 and 21, respectively (Extended Data Fig. 6d).

**Figure 5.**
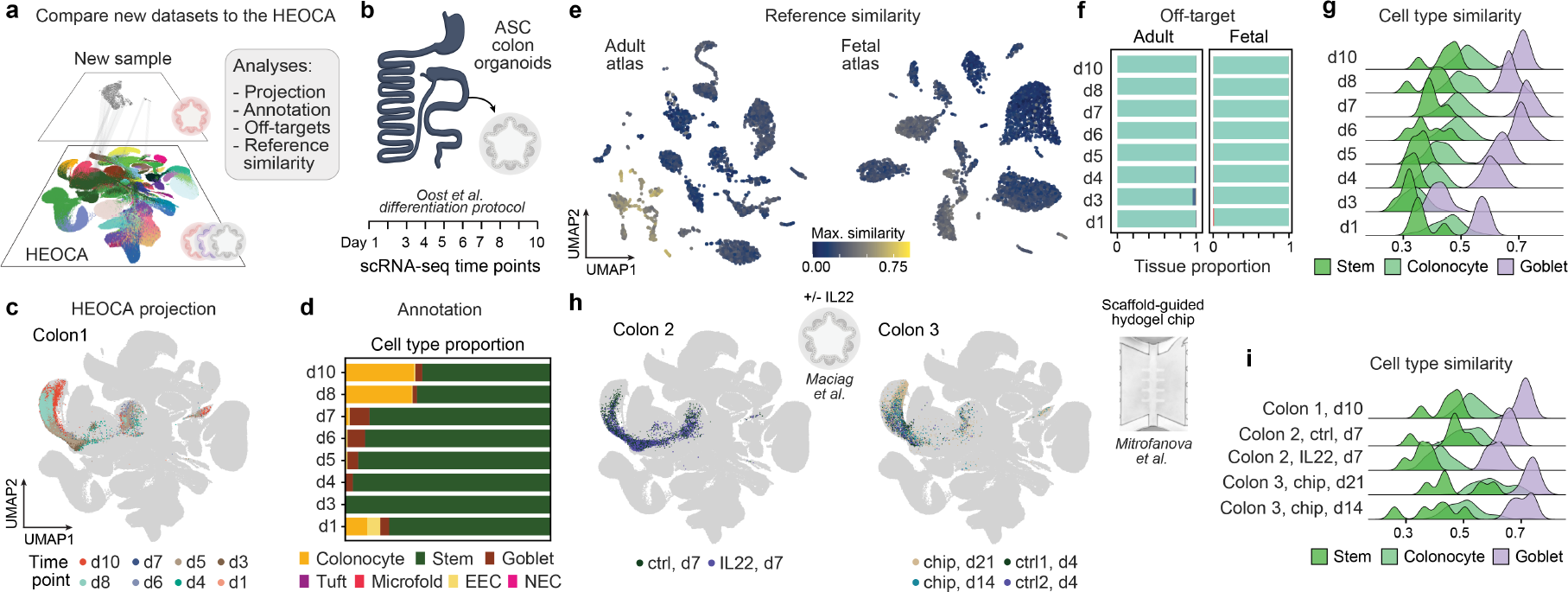
Assessment of new protocols and incorporation of new datasets into the HEOCA. (a) A summary describing the comparison of new datasets to HEOCA and subsequent analyses. (b) Experimental design of the new colon organoid time course samples. (c) UMAP of new colon protocol time course scRNA-seq data mapped to the organoid atlas colored by different time points. (d) The barplot depicts the cell proportions of predicted level 2 cell types across new colon protocol time course scRNA-seq data. (enteroendocrine cell (EEC), neuroendocrine cell (NEC)) (e) UMAP representations of adult (left) and fetal (right) primary tissues are displayed, highlighting the maximum similarity achieved by new organoid protocol within the comprehensive cross-tissue atlas. (f) The barplots showing the tissue proportion of the most similar adult (left) and fetal (right) tissue across new colon protocol time course samples. (g) Ridge plots display new colon protocol time course samples similar to adult colon primary tissue cell types. (h) Two additional scRNA-seq datasets from two new colon organoid protocols were integrated and mapped onto the organoid atlas. (i) Ridge plots display two new colon protocol samples similar to adult colon primary tissue cell types and compared to the time course day 10 sample.

Interestingly, this on-chip protocol has an advantage to conventional organoid protocols as it provides access to the apical and basal side of the epithelium and can be kept for many weeks in culture while maintaining stem and differentiated cell types.

Lung 1 and lung 2 datasets consisted of time courses of lung progenitor organoids differentiated to alveolar or airway organoids. The results demonstrated that an average of 99.27% and 95.56% of cells from these datasets were successfully integrated into the existing lung data within the atlas (Extended Data Fig. 6g-n). In the alveolar dataset, cells showed increased mapping to alveolar epithelial (AT1/AT2) identities over the course of differentiation. This was accompanied by a loss of cells mapping to undifferentiated identities in the reference, demonstrating that the reference atlas accurately assigned cells to alveolar epithelial identities and resolved the distinction between uncommitted progenitors and differentiating cells in the organoids. Likewise, lung progenitor organoids differentiated towards airway were accurately mapped to cell identities specific to the airways including SCGB3A2+ airway progenitors, basal and secretory cells, consistent with prior descriptions of these organoids (Conchola et al., 2023; Miller et al., 2020) and were minimally mapped to alveolar epithelial identities (Extended Data Fig. 6g-n). Altogether, these results highlight the utility of the HEOCA for integrating new data and assessing the effects of different experimental conditions or culture protocols on the maturation and cell type composition across diverse organoids.

### Mapping organoid disease models to the HEOCA illuminates disease-associated states

We next wanted to assess the utility of the integrated atlas to understand organoid models of disease. Through comparison to the HEOCA we assess cell proportion, identify disease-associated states, and perform differential expression analysis against the atlas data (Fig. 6). As a proof of principle, we first explored colorectal cancer (CRC) using a dataset composed of CRC organoids from a patient resection and normal organoids from adjacent healthy tissue (Wang et al., 2022)(Fig. 6a). After mapping to the HEOCA via scPoli, cell proportion analysis showed that CRC samples exhibited a significantly lower percentage of mature colonocytes, but a higher proportion of stem cells (Fig. 6b, Extended Data Fig. 7a-b). Interestingly, we also observed the emergence of mesothelial cells in the CRC samples, consistent with the published findings that CRC can lead to an increase in mesenchymal cells (Fig. 6b, Extended Data Fig. 7b) (Wang et al., 2022). We next calculated the mean distance of the top 100 nearest neighbors of each adjacent normal and cancer cell to the integrated atlas, and discovered that this distance metric could distinguish between normal and cancer cells (Fig. 6c, Extended Data Fig. 7c). Stem cells and colonocytes exhibited more pronounced deviation from atlas states, whereas goblet cells displayed fewer differences (Fig. 6d). We subsetted and integrated colonocytes from both adjacent normal organoid and cancer organoid samples, and discerned two distinct colonocyte groups (Fig. 6e). The first group comprised a mixture of normal and cancer organoid cells, while the second consisted exclusively of a subset of cancer organoid cells (Fig. 6e-f), which possessed a significantly larger distance to the HEOCA relative to the mixed group (Fig. 6g-h). Differential gene expression analysis revealed heightened expression levels of CRC markers such as CEACAM6, MMP7, TGFBI and RSPO3 in the cancer cell group (Fig. 6h, Extended Data Fig. 7d). Notably, recurrent R-spondin (RSPO) gene fusions have been described in certain CRC patients and this event potentiates Wnt signaling and tumorigenesis (Seshagiri et al., 2012). These analyses show the utility of distance measures to the HEOCA as a strategy to elucidate cell states that deviate healthy or otherwise normal states.

**Figure 6.**
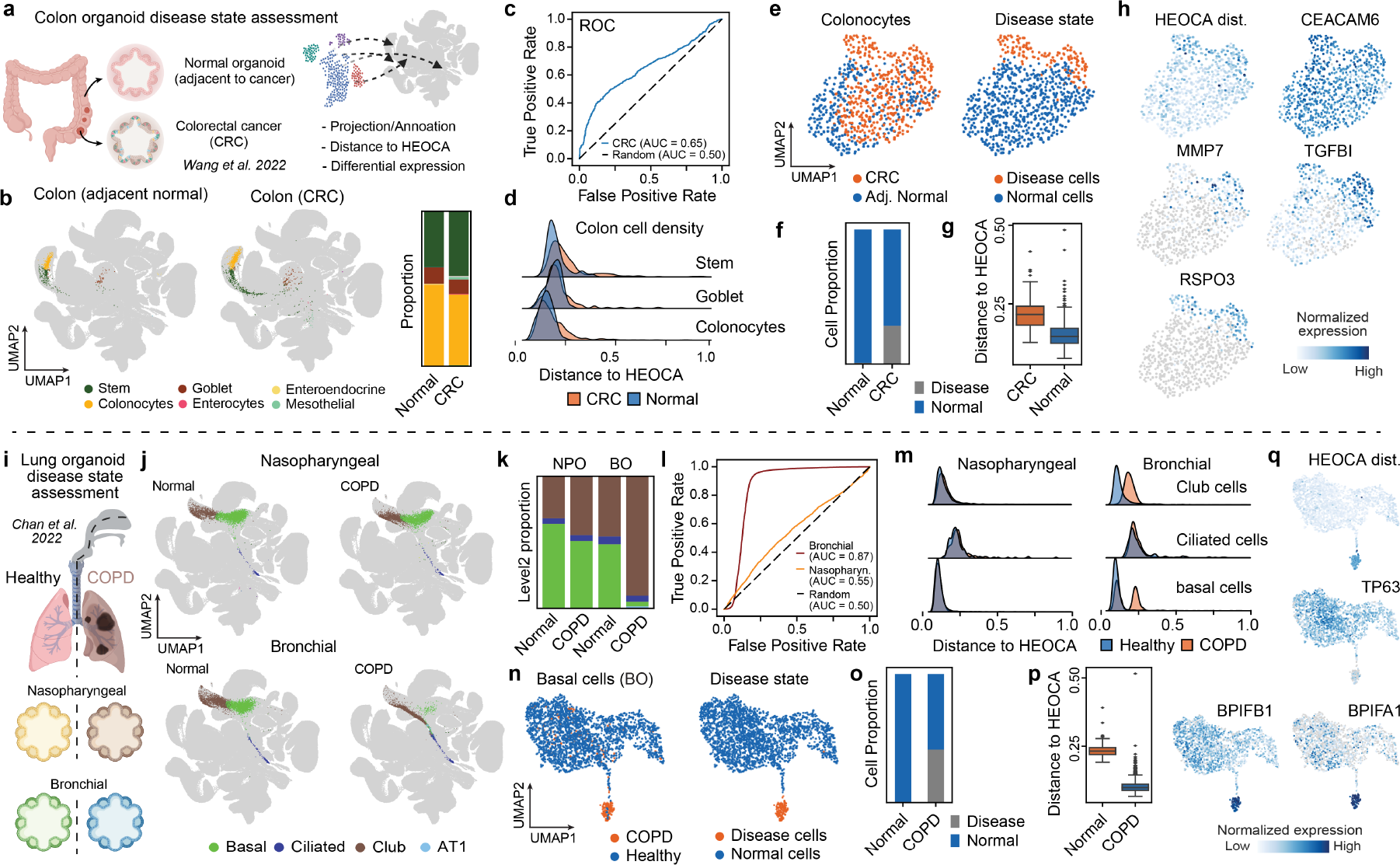
Comparison to the HEOCA reference of healthy organoids reveals disease-associated features. (a) An overview of colorectal cancer organoid samples and their analysis through the HEOCA assessment. (b) UMAP of normal colon and colorectal cancer scRNA-seq samples mapped to the HEOCA colored by predicted level 2 cell type. Cell proportion of predicted level 2 cell type in normal colon and colorectal cancer scRNA-seq samples. (c) ROC plot of prediction of colorectal cancer cells using the mean distance between single cells and their nearest neighbors in the organoid atlas. (d) The distance from adjacent normal and CRC cells to HEOCA is split by cell types. (e) UMAP plot of CRC and adjacent normal colonocytes, with cells color-coded based on their disease or normal sample (left) and their predicted disease state (right). (f) The barplot illustrates the distribution of CRC and adjacent normal cells within two distinct clusters of disease-state cells. (g) The boxplot presents the mean distance of the top 100 nearest neighbors to HEOCA for the two clusters of disease-state cells. (h) UMAP plot of CRC and adjacent normal colonocytes colored by the distance of cells to HEOCA and CRC marker gene expression. (i) An overview of COPD organoid samples and their analysis through the HEOCA assessment. (j) UMAP of normal nasopharyngeal, COPD nasopharyngeal, normal bronchial, and COPD bronchial scRNA-seq sample mapped to the HEOCA colored by predicted level 2 cell type. (k) Cell proportion of predicted level 2 cell type in normal nasopharyngeal, COPD nasopharyngeal, normal bronchial, and COPD bronchial scRNA-seq samples. (l) ROC plot of prediction of COPD cells using the mean distance between single cells and their nearest neighbors in the organoid atlas. (m) The distance from normal and COPD nasopharyngeal organoid (left) and normal and COPD bronchial organoid (right) to HEOCA is split by cell types. (n) UMAP plot of COPD bronchial and normal bronchial basal cells, with cells color-coded based on their disease or normal sample (left) and their predicted disease state (right). (o) The barplot illustrates the distribution of COPD bronchial and normal bronchial basal cells within two distinct clusters of disease-state cells. (p) The boxplot presents the distance of cells to HEOCA for the two clusters of disease-state cells. (q) UMAP plot of COPD bronchial and normal bronchial basal cells colored by the distance to HEOCA and COPD marker gene expression.

In a second assessment, we used a publicly available dataset of two different organoid types generated from cells of patients with chronic obstructive pulmonary disease (COPD) (Fig. 6i) (Chan et al., 2022). These were derived from nasopharyngeal and bronchial stem cells of these patients respectively. Intriguingly, both samples mapped to lung populations within the HEOCA, however there were substantial differences in the bronchial and nasopharyngeal mappings between normal and COPD conditions (Fig. 6j). Nasopharyngeal COPD organoids exhibited cell type composition similar to the healthy samples in the atlas, however bronchial organoids had a much larger proportion of annotated club cells and much fewer basal cells (Fig. 6j-k, Extended Data Fig. 7e-f). As was observed with the CRC organoids, distance to HEOCA states served as a strong metric to distinguish between normal and disease conditions, with a strong effect in the COPD-derived bronchial organoids (Fig. 6l, Extended Data Fig. 7g). These results matched with the original publication (Chan et al., 2022), which showed similar differences in cell type composition and reported differences in resistance to viral infection between the bronchial and nasopharyngeal COPD organoids. Based on atlas similarity, we observed that the nasopharyngeal normal and COPD samples showed relatively minor differences across all cell types, whereas basal cells in the bronchial COPD samples displayed a bimodal distribution (Fig. 6m). Distance to HEOCA states identified one basal cell population indicative of a disease state, which was further clarified in a heterogeneity analysis of basal cells from healthy and bronchial organoids (Fig. 6n-p). Differential expression analysis revealed decreased TP63 expression, and high expression of genes known to be upregulated in COPD such as BPIFA1 and BPIFB1 (Fig. 6q, Extended Data Fig. 7h). Together these data show that the HEOCA can be used to place cell states observed in organoid disease models into a larger context, which helps to better understand holistic effects on cell composition and gene expression patterns.

## Discussion

Single-cell transcriptome sequencing technologies have advanced organoid research by offering a powerful set of experimental and computational tools to investigate cell types present in these complex 3D models. Despite the immense progress, it remains a challenge to understand and quantify organoid fidelity and to place variation between organoid datasets into a larger context of human physiology. To begin to address these challenges, we have built the first integrated cell atlas of organoids that model endoderm-derived tissues, incorporating organoid datasets that have been generated from multiple different types of stem cells and protocols. We have established a framework for integration and harmonized cell type annotation, which makes interpreting cell heterogeneity between organoid datasets tractable. Single-cell transcriptome data from diverse experimental designs can introduce strong technical noise due to batch effects, protocol variation, genomic method, and other technical biases, making data integration challenging. To overcome this, we evaluated ten existing integration methods and identified an approach with excellent performance. This integration method, scPoli, is structured to incorporate newly generated data, enabling rapid comparison of novel datasets across the nearly 1 million organoid cells in the current atlas. Consolidation and integration of such a diversity of organoid cell states provides new inroads into quantifying fidelity. Indeed, we find that there is significant variation in organoid cell composition, prevalence of off-target cells, and overall cell state similarity. This variation, and comparison to available reference atlases, revealed that current organoid technologies cover a large diversity of human cell types and states. This result clarifies the use of human organoid technologies to explore development, model disease, and identify therapeutics.

Through cross-organ, multi-organoid integration, it was possible to identify off-target cells, a particular problem in PSC-derived organoids due to incomplete specification, as well as to distinguish cell states that significantly differed from states present in the atlas. This ability to distinguish non-present states is helpful to assess new protocols, as well as to identify features of disease models that are non-variable in normal, healthy organoids. Indeed, the integrated HEOCA presents a first opportunity to place a new endodermal organoid dataset into a relationship with the accumulated other datasets generated to date. There is still a major challenge with organoid fidelity quantification, as there is not yet a complete and integrated atlas of human cell type diversity during development and adulthood from primary tissues. Comprehensive integrated reference atlases across the human lifespan, in health and disease, together with diverse organoid models in normal and perturbed conditions, will help to clarify the full potential of the human genome. Altogether, the HEOCA will serve as a valuable resource for the organoid research community and a foundation to expand the ability to model human biology.

## Data availability

The HEOCA (raw and normalized counts, integrated embedding, cell type annotations, and technical metadata) data will be made available in CELLxGENE Discover when the final paper is accepted. The HEOCA core reference model and embedding for the mapping of new data to the HEOCA and human intestinal organoid cell atlas can moreover be found on Zenodo (https://doi.org/10.5281/zenodo.8181495).

## Code availability and analytic reproducibility

Code for scRNA-seq cell type annotation is available at GitHub (https://github.com/devsystemslab/snapseed), and code for mapping news data and disease data to HEOCA is available at GitHub (https://github.com/devsystemslab/sc2heoca), code for differential expression gene analysis is available at GitHub (https://github.com/bioqxu/wilcoxauc), code for all the analysis in this paper is available at GitHub (https://github.com/devsystemslab/HEOCA).

## Author contributions

Q.X. downloaded the data, integrated the HEOCA and intestinal organoid cell atlas and performed downstream analysis of HEOCA and intestinal atlas. L.H., S.H., and M.K. were involved in the integration of the lung organoid cell atlas and the downstream analysis of HEOCA, intestinal and lung organoid cell atlas. T.F. provided expertise in lung organoid development and assisted in re-annotation and harmonization of organoid and primary lung tissue. U.K., J.B. performed the new organoid single cell RNA-seq in this paper and edited the manuscript. S.P., M.G., R.K.F. and D.K. performed experiments in the analysis of the lung and intestinal atlas. Q.Y., L.A., Z.H., and J.S.F. helped integrate the atlas and edited the manuscript. K.O., M.K., S.B., O.M., G.M., K.B.J., M.L., P.L., J.R.S. provided datasets and edited the manuscript. Q.X., L.H., B.T., F.T., and J.G.C. wrote the manuscript. B.T., F.T., and J.G.C. conceived the study and supervised analysis of the integrated data. All authors reviewed the manuscript.

## Acknowledgements

This publication is part of the Human Cell Atlas (www.humancellatlas.org/publications/). We thank T. Gomes and R. Okuda for the helpful discussions on the cell markers of organoid cell types. We are grateful to Julien Gagneur and his IT team for access to GagneurLab computing resources for the duration of study. We extend our gratitude to the CELLxGENE team (https://cellxgene.cziscience.com/) for their invaluable assistance in organizing and facilitating the publication of our atlas as a cell browser. J.G.C., J.R.S., and B.T. are supported by grant CZF2019-002440 from the Chan Zuckerberg Initiative DAF, an advised fund of the Silicon Valley Community Foundation. The Novo Nordisk Foundation Center for Stem Cell Medicine is supported by a Novo Nordisk Foundation grant (NNF21CC0073729). J.G.C. is supported by the European Research Council (Anthropoid-803441). This project has received funding from the European Union’s Horizon 2020 research and innovation programme under grant agreement No 874769 and the Chan Zuckerberg Foundation Grant CZF 2019-002440 to P.L.. The work on bioengineered human mini-colons was funded by support from the Swiss National Science Foundation (SNSF) research grant 310030_179447, the EU Horizon 2020 Project INTENS (#668294-2) and Ecole Polytechnique Fédérale de Lausanne (EPFL). F.T. is co-funded by the European Union (ERC, DeepCell – 101054957). Views and opinions expressed are however those of the authors only and do not necessarily reflect those of the European Union or the European Research Council. Neither the European Union nor the granting authority can be held responsible for them.

## Methods

### Generation and culture of PSC-derived human intestinal organoids

All PSC lines were cultured with mTESR/mTESR-plus (STEMCELL Technologies 85850/ 100-0276) media on Matrigel (Corning, 354277) coated plates. The cells were passaged when they reached 80-90% confluency every 4-5 days. PSCs were differentiated into HIO following the previously described protocols (Spence et al., 2011; Tsai et al., 2016; Capeling et al., 2020). In brief, PSCs were induced into definitive endoderm by using 100 ng/ml Activin A (R&D, 338-AC) in RPMI-1640 for three days with increasing concentrations of FBS (0%, 0.2%, and 2%). Then, midgut/hindgut patterning of endoderm was directed with 500 ng/ml FGF4 (Peprotech, 100-31) and 2uM CHIR99021 (Stem Cell Technologies, 72052) with daily media changes for up to 6 days in the presence of 2% fetal bovine serum (FBS). Spheroids were collected after 96, 120, and 144 hours of patterning and embedded in Matrigel (Corning, 354234/356231), cultured in ENR media (mini gut basal media supplemented with 100 ng/ml EGF (R&D, 236-EG), 100 ng/ml purified Noggin-Fc (Heijmans et al., 2013) and R-Spondin-1-Fc (5% conditioned media) (Ootani et al., 2009). The mini gut basal media consists of DMEM/F-12, HEPES (Thermo Fisher Scientific, 11330032), and 1x B27 supplement (Thermo Scientific, 12587001). All media used in the differentiation process contain 1x Penicillin-Streptomycin (Thermo Scientific, 15140122). Organoid media was changed every 3 days. HIOs were split once the mesenchyme outgrew Matrigel, and excessive cell debris accumulated in the core every 7-10 days.

For the suspension HIO protocol (Capeling et al., 2022), the modifications to the original protocol described above is that the budded spheroids were collected and cultured in ultra-low attachment plates without splitting to protect the mesothelial layer. During organoid growth, Noggin and R-spondin were removed after 3 days of growth.

Another exception to the main protocol was HIO-ECs (vHIO) (Holloway et al., 2020) grown with minor adjustments to introduce endothelial cells co-development with a first three days of spheroid growth with minigut media containing EGF (100 ng/ml), VEGF (50 ng/ml), bFGF (25 ng/ml), and BMP4 (25 ng/ml) that followed by supplementation with EGF (100 ng/ml) and VEGF (25 ng/ml) for the rest of the in vitro growth of the organoids as described previously.

### Human intestinal organoid transplantation

HIOs were cultured in Matrigel domes in ENR media for a period of 4 weeks until transplantation. All the HIOs were grown in ENR media in vitro except the iPSC72.3- and two H9-derived HIOs which were grown with ENR media till day 3 and then changed to EGF media (Holloway et al., 2020) and passaged weekly. On the day of transplantation, HIOs were mechanically dissociated from the Matrigel and implanted beneath the kidney capsules of immunocompromised NOD-SCID IL2Rg null (NSG) mice (Jackson Laboratory, strain number 0005557), following established protocols (Watson et al., 2014; Finkbeiner et al., 2015). Briefly, mice were anesthetized using 2% isoflurane, and a left-flank incision was made to expose the kidney after proper shaving and sterilization with isopropyl alcohol. HIOs were implanted beneath the mouse kidney capsules using forceps. Prior to closure, an intraperitoneal flush of Zosyn (100 mg/kg; Pfizer) was administered. Mice were euthanized after 8 weeks for scRNA-seq. For particular experiments, location of transplantation was changed to mesentery (Cortez et al., 2018).

### Media compositions for human enteroids

To support growth and maintenance of human ASC-derived intestinal organoids in 3D we culture organoids in Matrigel domes, WERN-complete media was used (Gu et al., 2022; Sato et al., 2011; Miyoshi and Stappenbeck, 2013). The composition of the complete media: 50% L-WRN Condition Medium (University of Michigan), GlutaMAX Supplement (2 mM, Thermo Fisher Scientific, Cat 35050061), HEPES (10 mM, Fisher, Cat 15630080), Primocin (100 μg/mL, Invivogen, Cat ant-pm-1), N-2 supplement (1×, Thermo Fisher Scientific, Cat 17502048), B-27 supplement (1×, Thermo Fisher Scientific, 17504044), N-Acetyl-L-Cysteine (1 mM, Sigma-Aldrich, A9165), Recombinant Human EGF Protein, CF (100 ng/mL, RD Systems, 236-EG-01M), A 83-01 (500 nM, Stem Cell Tech, 72022), SB 202190 (100 μM, Stem Cell Technologies, 72632), Nicotinamide (10 nM, Sigma-Aldrich, Cat N5535), and Advanced DMEM/F-12 (Thermo Fisher Scientific, Cat 12634028).

The composition of the budding differentiation media for human enteroids adapted (Fujii et al., 2015) with minor modifications : 50% L-WRN Condition Medium (University of Michigan), GlutaMAX Supplement (2 mM, Thermo Fisher Scientific, Cat 35050061), HEPES (10 mM, Fisher, Cat 15630080), Primocin (100 μg/mL, Invivogen, Cat ant-pm-1), N-2 supplement (1×, Thermo Fisher Scientific, Cat 17502048), B-27 supplement (1×, Thermo Fisher Scientific, 17504044), N-Acetyl-L-Cysteine (1 mM, Sigma-Aldrich, A9165), IGF1 (100 ng/mL, BioLegend, Cat 590904), FGF-basic (50, ng/mLPeprotech, Cat 100-18B), and A 83-01 (500 nM, Stem Cell Tech, 72022), Nicotinamide (10 nM, Sigma-Aldrich, Cat N5535), and Advanced DMEM/F-12 (Thermo Fisher Scientific, Cat 12634028).

### Single-cell dissociation of organoids and tissues

PSC-derived human organoids, tHIOs and newly generated ASC-derived enteroids dissociated to single cells using the previously established protocol (Yu et al., 2021). Briefly, to prevent cell adhesion, all tubes and pipette tips were coated with 1% BSA in HBSS. The organoids were mechanically dislodged, pipetted up and down to remove excess Matrigel, and pooled in a Petri dish. After removing excess media, the organoids were minced into smaller fragments in HBSS. The tissue pieces were transferred to a 5 ml conical tube containing Mix 1 from the Neural Tissue Dissociation Kit (Miltenyi Biotec, 130-092-628) and incubated on a shaker for 15 minutes at room temperature. Mix 2 was added to the tube, and every 10 minutes for another 20-30 minutes, the mix was agitated with a P1000 pipette tip. The cells were then filtered through a 70 μm strainer and washed three times at 400g for 5 minutes with 1% BSA in HBSS. Cell counting was performed using Countess and the process was immediately continued with 10x Chromium for single-cell droplet generation. The libraries were prepared using the NextGEM Single Cell 3’ v3.1 kit according to the manufacturer’s instructions. The libraries were sequenced on Illumina’s Novaseq6000 system. For detailed information on each library and associated metadata refer to the Supplementary Table.

### Human liver organoid

Human fetal liver tissue (16 gestational week) was obtained following elective pregnancy termination and informed written maternal consents from Cercle Allocation Services, USA. Tissue dissection and organoid culture were performed as described previously (Hu et al., 2018). The organoid culture at the moment of characterization contained a mixture of phenotypic hepatocyte- and cholangiocyte-enriched organoids.

### Human stomach organoid

Human stomach tissue was obtained and experimental procedures performed within the framework of the non-profit foundation HTCR (Munich, Germany) including informed patient’s consent. Healthy tissue was obtained as part of resections on gastric cancer, and both antrum and pylorus organoids were established from the same donor. Tissue dissection and organoid culture were performed as described previously (Bartfeld et al., 2015).

### Time course colon organoids

Mature organoids are collected by rinsing the maintenance well using 1-2 ml cold DMEM/F12 (Stem Cell Technologies) + 15 mM HEPES + 1x GlutaMax (1:100 35050-038) + 100 μg/ml PenStrep (15140-122) (=DMEM+++) and transferring the solution into a 15 ml falcon tube. Next, DMEM+++ is added until a total volume of 7 ml is reached before centrifuging the tube at 400 rcf for 4 min at 4°C. Supernatant is discarded and organoids are resuspended into 2 ml of prewarmed 0.05% Trypsin + EDTA (gibco, 25300-054) and incubated for 10 min at 37°C. Trypsinization is stopped using 2 ml of a trypsin-inhibitor (Thermo Fisher Scientific, 17075029) as well as 3 ml of DMEM+++ after reaching a single-cell suspension. Singularized organoids are then centrifuged at 600 rcf for 5 min at 4°C and supernatant is discarded subsequently. Next, the cell pellet is resuspended into START medium and then filtered through a 40 μm filter to remove large debris and cell clumps. Optional: FAC-sorting for single cells to remove remaining smaller debris as well as doublets. Cells are then counted before being further processed. Finally, the cell-suspension is diluted to 25-50 cells/μL into a 60% Matrigel : 40% START medium mixture (we have tested dilutions with a Matrigel proportion of 50 to 70%). Here, lower seeding densities between 25 and 50 cells per μL are recommended as very dense plating affects the morphology and growth of organoids negatively – this needs to be optimized for every line. For maintenance, the Matrigel mixture is plated into cell culture plates in 5-10 μL droplets and are covered after a 30 min 37°C solidification period with warm START medium. Medium is changed after 4 days to BALANCE, and is refreshed once with new BALANCE medium at day 7 until maturation is reached, typically at day 10. START and BALANCE media compositions are described in Oost et al., 2023 (currently in revision and will be made available).

### Mini-colon organoid

Human colon organoids were maintained for 4 days starting from mechanically dissociated fragments in Matrigel (Corning, 356231) or Matrigel supplemented with 20% Collagen I (Advanced Biomatrix, 5225) in Advanced DMEM/F12 with penicillin/streptomycin, 1× Glutamax and 10 mM HEPES, supplemented with 1× B27 supplement (Gibco), 1 μM N-acetylcysteine (Sigma-Aldrich), 0.5 nM Wnt Surrogate-Fc Fusion Protein (U-Protein Express B.V., N001), 100 ng/ml Noggin (EPFL Protein Expression Core Facility), 500 ng/ml R-Spondin 1 (EPFL Protein Expression Core Facility), 100 ng/ml recombinant human IGF-1 (BioLegend), 50 ng/ml recombinant human FGF-2 (Peprotech), 10 nM gastrin, 100 ng/ml recombinant human NRG1 (RD, 5898-NR) and 500 nM A83-01 (Tocris). Human mini-colons were maintained for 14 and 21 days, where starting from day 7 apical medium was switched to ‘ENR’ composed of 50 ng/ml recombinant human EGF (RD, 236-EG), 100 ng/ml Noggin (EPFL Protein Expression Core Facility), 500 ng/ml R-Spondin 1 (EPFL Protein Expression Core Facility), and organoid medium as described above was maintained on the basal side.

Human mini-colons cultured in biological triplicates were extracted from the PDMS chips and digested with collagenase I (Sigma) at 100 U/ml concentration in Advanced DMEM/F12 with penicillin/streptomycin, 1× Glutamax and 10 mM HEPES, supplemented with 1× B27 supplement (Gibco), 1 μM N-acetylcysteine (Sigma) for 10 min at 37°C with 300 rpm. Human colon organoids in biological triplicates were collected from the Matrigel domes, and both organoids and mini-colons were dissociated to single cells using 4 mg/mL Protease VIII (Sigma) in dPBS containing 10 μM Y-27632 on ice for approximately 45 min with trituration using a P1000 micropipette every 10 minutes. Following one washing step in PBS+2% BSA (Gibco), cell suspensions were incubated for 20 min on ice with TotalSeq™-C anti-human Hashtag oligos (HTOs) (1:500, Biolegend, 394661, 394663, 394665, 394667, 394669, 394671, 394673, 394675, 394677, 394679, 394683, 394685) in PBS+2% BSA. Cells from each condition and replicate were washed three times with PBS+2% BSA, pooled together and filtered through low-volume 10 μm cell strainers (PluriSelect). All cell suspensions were recounted to achieve a uniform concentration of 2000 cells per microliter before pooling for 10× capture. The cell hashing and cDNA libraries were constructed using 10x Genomics Chromium Next GEM Single Cell 5’ Reagent Kits v2 (Dual Index) reagents and sequenced using Illumina protocol using NovaSeq 6000 reagents with around 50000 reads per cell.

### IL22-/+ colon organoid

Human colonic epithelial organoids from three healthy individuals were cultured using previously published culture conditions (Fujii et al., 2015). Organoids were seeded as single cells and treated with IL-22 (10 ng/mL) from day 4 to day 7. Treated and untreated organoids were collected at day 7 and dissociated to single cells using TrypLE (Invitrogen) incubation at 37°C. The six different samples were tagged with different TotalSeqTM hashtag antibodies (A0251-A0256, Biolegend). Cells were subsequently sorted on an ARIAII FACS sorter (BD) to isolated DAPI negative cells. A total of 6100 events/cells were isolated from each sample for scRNA-sequencing using the 10X Genomics protocol v3.1 with implementation of hashtag libraries. Additional HTO primer (0.2 μM) was added in the step of cDNA amplification to increase yield of hashtag-barcodes. After the cDNA amplification, hashtag-cDNA and endogenous cDNA were separated based on the size with SPRIselect beads (Beckman Coulter) as described (Stoeckius et al., 2018). The dual index kit set AA (10X genomics) were used for sample indexing and 11 cycles were used during the final amplification step. The hashtag cDNA and endogenous cDNA libraries were diluted to 4 nM and pooled (5% hashtag + 95% endogenous cDNA) before being sequenced on a NextSeq2000 sequencer (Illumina).

### Data collection

The scRNA-seq data used in this study were obtained from the original papers (Extended Data Table 1). If the raw fastq files were available, they were downloaded. The seq2science method (https://github.com/vanheeringen-lab/seq2science) was used to download the raw fastq files in Gene Expression Omnibus (GEO) database (https://www.ncbi.nlm.nih.gov/geo/) or BioStudies database (https://www.ebi.ac.uk/biostudies/). The reads were aligned to the GRCh38 genome and Ensembl 98 gene annotation using the STARsolo (Kaminow et al., 2021). In cases where the raw FASTQ files were not available, the raw counts were downloaded instead. The downloaded counts and the counts obtained from the realigned reads were merged for subsequent analysis.

### Data normalization

To integrate the data, we combined the count data from all the samples into a unified dataset. For subsequent analysis, we retained only the genes classified as protein-coding genes and long non-coding RNA (lncRNA) genes. The low quality cells in each sample were filed. The raw counts were then normalized to a total count of 10,000 and log-transformed. Given these normalized counts, the top 3,000 highly variable genes were identified using the default settings in Scanpy. These highly variable genes were selected for further downstream analysis.

### Cell type annotation

For each sample, the raw counts were normalized to a total count of 10,000 and then log-transformed. From these normalized counts, the top 3,000 highly variable genes were identified using the default settings in Scanpy. These highly variable genes were selected as the subset for further downstream analysis. Principal component analysis (PCA) was performed on the normalized data, and the top 30 principal components were chosen for calculating the K Nearest Neighbors (KNNs). Using the KNNs, a Uniform Manifold Approximation and Projection (UMAP) was generated to visualize the data in a lower-dimensional space. To cluster the data, the Leiden clustering method with a resolution of 2 was applied. This clustering approach helped to identify distinct groups of cells based on their gene expression patterns. Previously defined marker genes associated with specific cell types were used to guide the annotation process. To annotate cell types within each cluster, the snapseed method was employed. This method calculates the area under the receiver operating characteristic (ROC) curve (AUC) and fold change values for each marker gene in relation to the cluster. If multiple markers were available for a particular cell type, the maximum AUC and fold change values were selected. The average AUC and fold change values were used to represent the specific cell type, and the most specific cell type was annotated for each cluster based on these criteria.

### Data integration benchmarking

To benchmark and compare different integration methods, 10 random samples were selected from the dataset. Several integration methods, including scVI, scANVI, scPoli, bbknn, harmony, combat, CSS (pearson), and CSS (spearman) (Polański et al., 2020; Xu et al., 2021; Lopez et al., 2018; Korsunsky et al., 2019; He et al., 2020; Büttner et al., 2019; De Donno et al., 2023), were applied to the data to assess their performance in integrating the samples. The scIB method, a benchmarking tool, was used to evaluate and compare the results obtained from these integration methods. In the scPoli model, we configured the following parameters for effective training and integration: embedding_dim was set to 3; hidden_layer_sizes were determined as the square root of the total number of cells. During the training phase, we employed the following settings: early_stopping_metric was set to val_prototype_loss; mode was set to min; threshold was set to 0; patience was set to 20; reduce_lr was enabled, with lr_patience set to 13 and lr_factor set to 0.1; n_epochs were set to 5; pretraining_epochs were set to 4; eta was set to 10; alpha_epoch_anneal was set to 100.

### Cell type reannotation

After integration, we re-cluster all cells in the atlas based on the scPoli integrated embedding using the Leiden method with a resolution of 10 (HEOCA and HIOCA) and 5 (HLOCA), respectively. New annotations were then assigned to each cluster using the dominant cell type per cluster.

### Marker gene refinement

We randomly subset 100K cells from the atlas. For each cell type, we used the wilcoxauc method (https://github.com/bioqxu/wilcoxauc) to calculate the Area Under the Curve (AUC), fold change, and adjusted p-value for each gene. Genes in each cell type were filtered, keeping those with AUC >= 0.6, log fold change >= 0.25, and adjusted p-value <= 0.01. These filtered genes were then ranked based on their AUC score, and the top 10 genes were selected as marker genes for each cell type. We combined the selected marker genes and performed hierarchical clustering on the resulting gene set.

### Cross-organ primary tissue integration

The 1 million subsets of the human fetal atlas were downloaded (https://cellxgene.cziscience.com/collections/c114c20f-1ef4-49a5-9c2e-d965787fb90c) (Cao et al., 2020). The normal endoderm tissues including the pancreas, lung, liver, intestine, and stomach were subsetted. The top 3000 highly variable genes were subsetted for data integration. The cells within each tissue were integrated using the scPoli method20, with the cell_type serving as the cell type key for integration, and with the same parameters used in the HEOCA atlas integration. The scPoli model was saved for the downstream comparison. The tabula sapiens multiple-organ adult single-cell transcriptomic atlas of humans were downloaded (https://tabula-sapiens-portal.ds.czbiohub.org/) (Tabula Sapiens Consortium* et al., 2022). The endoderm tissues including the liver, lung, pancreas, small intestine, large intestine, prostate and stomach were subsetted. The endothelial, epithelial, and stromal compartment of cells were subsetted. The top 3000 highly variable genes were subsetted for data integration. The cells within each tissue were integrated using the scPoli method (De Donno et al., 2023), with the cell_ontology_class serving as the cell type key for integration, and with the same parameters used in the HEOCA atlas integration. The scPoli model was saved for the downstream comparison.

### Correlation to primary tissue

To compare and correlate cell states in primary tissue and organoid models, miloR (Dann et al., 2022) method was used to define and construct neighborhood graphs for each data source separately. We computed the transcriptional similarity graph for the primary tissue reference using 30 nearest neighbors and the UMAP representation of latent representations of integrated primary tissue cells. To compute the transcriptional similarity graph for the organoid reference, we used the 30 nearest neighbors and the UMAP representation of integrated embedding of organoid cells. Single cell organoid data were integrated using scPoli and 3000 highly variable genes as described earlier. We used the default parameters for all the remaining computational steps in building the neighborhood graphs. We then used the R package scrabbitr (Ton et al., 2023) to compute the correlation between each pair of neighborhoods in the primary tissue and organoid reference and annotate the results at cell type or tissue level. The neighborhood correlations were computed using 3000 highly variable genes that were found in the highly variable genes in the primary tissue single cell reference atlases. This step results in two neighborhood correlation matrices: a primary tissue-correlation matrix where each entry marks the correlation of the expression profile of a given neighborhood in the primary tissue with the HEOCA, and an organoid-correlation matrix that stores the correlation of expression profiles in each neighborhood of the organoid atlas with the primary tissue atlas. This procedure was additionally repeated for each organoid derivation protocol, that is Adult Stem-Cell, Fetal Stem-Cell and Pluripotent Stem-Cell derived protocols. To compare the correlation between cell states in the primary tissue and between organoid derivation protocols, we subtracted the primary tissue-neighborhood correlation matrices computed with respect to neighborhoods for each derivation protocol. This approach of comparing primary tissue and organoid by correlation of neighborhood graphs is more reliable than the alternative reference mapping strategy, as it removes the dependance of the reliability and accuracy of the conclusions to mapping uncertainty, and allows for computing correlation statistics on graphs that are constructed based on transcriptional similarity of cells in each data source.

### Organoid off-target analysis

For each organoid sample, the same set of variable genes used in the primary tissue atlas (adult or fetal) was chosen, and the scPoli query was executed using identical parameters to those employed in the primary tissue atlas training model. The UMAP embedding was transformed using the primary tissue atlas UMAP model. For each new cell, the system selected its 100 nearest neighbors from the HEOCA dataset. The predicted tissue for the cell was assigned based on the tissue that was most frequently observed among its 100 nearest neighbors.

### Velocity and pseudotime analysis

For RNA velocity analysis of the HIOCA we first excluded samples missing splicing information. We then applied scVelo (Bergen et al., 2020) to generate a UMAP representation with stream trajectory visualization. The velocity pseudotime, spanning from stem cells to enterocytes and colonocytes, has been rescaled to a range of 0 to 1. We calculated and displayed the average expression of markers within specific bins.

For trajectory associated gene analysis, we first applied scanpy.tl.dpt diffusion pseudotime trajectory inference as described before with a stem cell set as the root cell (Haghverdi et al., 2016). Based on this, we generated stream plots for trajectory visualization. We then used CellRank (Lange et al., 2022) to determine macrostates and chose at least one terminal state per included cell type. CellRank was subsequently used to uncover putative driver genes by correlating fate probabilities with gene expression. Identified driver genes were plotted as waterfall plots per cell type ordered by the respective gene expression peaks in pseudotime.

### Intestine organoid atlas integration

The top 3000 highly variable genes were subsetted for data integration. To integrate all the cells, we applied the scPoli method with the same parameters used in the HEOCA atlas integration. The scPoli model was saved for the downstream comparison.

### Lung organoid atlas integration

Lung organoid single cell data curated from different studies was subsetted on top 3000 highly variable genes for integration. We applied scPoli to learn 30-dimensional latent representations of the cells, and 10-dimensional latent representations of the samples using a neural network with 2 hidden layers each of size 512. The network was trained setting pre-training epochs to 4, eta=10, patience=20, lr_patience=13, lr_factor=0.1, alpha_epoch_-anneal=100, reduced_lr=True and prototypical loss of the validation set as the early stopping criteria. The scPoli model was saved for the downstream comparison.

### Intestine primary tissue atlas integration and compression of organoid samples

The scRNA-seq data from both duodenum fetal and adult primary tissues were obtained from two research papers (Yu et al., 2021; Elmentaite et al., 2021). We focused on epithelial cells and subsetted them for analysis. The top 3000 highly variable genes were subsetted for data integration. To integrate all the cells, we applied the scPoli method with the same parameters used in the HEOCA atlas integration. The scPoli model was saved for the downstream comparison. For each organoid sample, the same set of variable genes used in the primary tissue atlas was chosen, and the scPoli query was executed using identical parameters to those employed in the primary tissue atlas training model. The UMAP embedding was transformed using the primary tissue atlas UMAP model. For each new cell, the system selected its 100 nearest neighbors from the primary tissue dataset. The predicted cell type for the cell was assigned based on the tissue that was most frequently observed among its 100 nearest neighbors.

### Lung primary tissue atlas integration and compression of organoid samples

Fetal data was obtained from the human fetal lung atlas (He et al., 2022) (https://fetal-lung.cellgeni.sanger.ac.uk/) and subset to epithelial cells. We provide original labels and manually simplified reannotation based on expert annotation. For mature data we obtained a dataset of similar size (GSE143868) (Deprez et al., 2020) included in the HLCA core reference (Sikkema et al., 2023) and annotated the cells using the Human_Lung_Atlas.pkl model implemented in the CellTypist annotation tool (Domínguez Conde et al., 2022). Data was subset to epithelial cells and the CellTypist-provided annotations were simplified by grouping fine-grained cell types into higher-order families. Organoid data was integrated and mapped onto the concatenated references using scPoli. The integration pipeline was run with raw counts and the top 3000 highly variable genes in the reference data. We subsequently followed the query-to-reference mapping workflow outlined in the scArches documentation (Lotfollahi et al., 2022). Once the query cells were mapped to the reference, predicted cell types and maturity were inferred based on the labels of the 10 nearest neighbor cells in the reference embedding. UMAP visualizations depict the reference embedding obtained from the trained query-to-reference model.

### Differential abundance analysis in the lung subset of HEOCA

We used Milo (Dann et al., 2022) to carry out differential abundance analysis between different stem-cell derived organoid lung samples in HEOCA to compare cell states between protocols. Milo neighborhood graph was constructed using the 2-dimensional UMAP representation of the lung data in HECOA integrated embedding, setting the k-nearest neighbor parameter to 30. We used default values for all other remaining parameters. Cell counts were quantified in each neighborhood and differential abundance (DA) was determined by fitting a generalized log-linear negative binomial model per neighborhood. Differential abundance calls were made at Spatial FDR < 0.05.

### New dataset incorporation

The new samples of scRNA-seq raw reads were mapped to the human genome, and counts of the matrix were obtained. The same set of variable genes used in HEOCA was chosen, and the scPoli (De Donno et al., 2023) query was executed using identical parameters to those employed in the HEOCA training model. The UMAP embedding was transformed using the HEOCA UMAP model. For each new cell, the system selected its 100 nearest neighbors from the HEOCA dataset. The predicted cell type for the new cell was determined by assigning it the cell type that was most frequently observed among its 100 nearest neighbors at the level 2 cell type classification level. Similarly, the predicted tissue for the new cell was assigned based on the tissue that was most frequently observed among its 100 nearest neighbors.

### Disease sample analysis

The disease sample analysis is similar to the new sample incorporation step. The raw count matrices were downloaded from the original papers. The same set of variable genes used in HEOCA was chosen, and the scPoli (De Donno et al., 2023) query was executed using identical parameters to those employed in the HEOCA training model. The UMAP embedding was transformed using the HEOCA UMAP model. For each new cell, the system selected its 100 nearest neighbors from the HEOCA dataset. The predicted cell type for the new cell was determined by assigning it the cell type that was most frequently observed among its 100 nearest neighbors at the level 2 cell type classification level. The predicted tissue for the new cell was assigned based on the tissue that was most frequently observed among its 100 nearest neighbors. For each cell, the mean distance of its 100 nearest neighbors was assigned as its mean distance to HEOCA.

In the analysis of differentially expressed genes between colon cancer organoid colonocytes and bronchial COPD organoid basal cells, we performed separate subsetting for all colonocytes and basal cells. For each dataset, we isolated the top 3000 highly variable genes. We then integrated these subsets of cells using the bbknn method (Polański et al., 2020). To cluster the two datasets, we applied the Leiden method with resolutions of 1 and 2 in two datasets. The clusters predominantly associated with the disease were selected as disease state cells, while the remaining clusters were categorized as normal state cells.

## Supplementary Information

**Figure S1.**
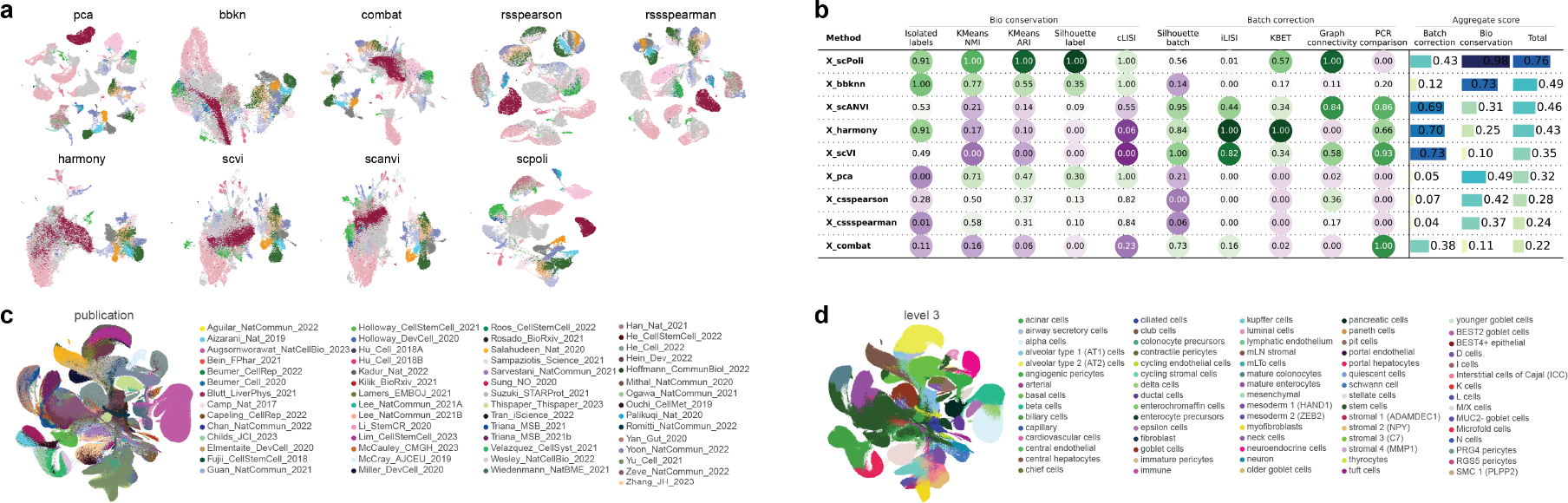
Analysis of scRNA-seq integration methods. (a) UMAP of tested integration methods and without any data integration (PCA). Dots in all UMAP embeddings are colored by the level 2 cell type annotation. (b) scIB benchmarking metrics for all tested integration methods. (c-d) UMAP of the organoid atlas colored by publications (c) and level 3 cell annotations (d). This is a supplementary figure shown as a two-column image with a legend underneath.

**Figure S2.**
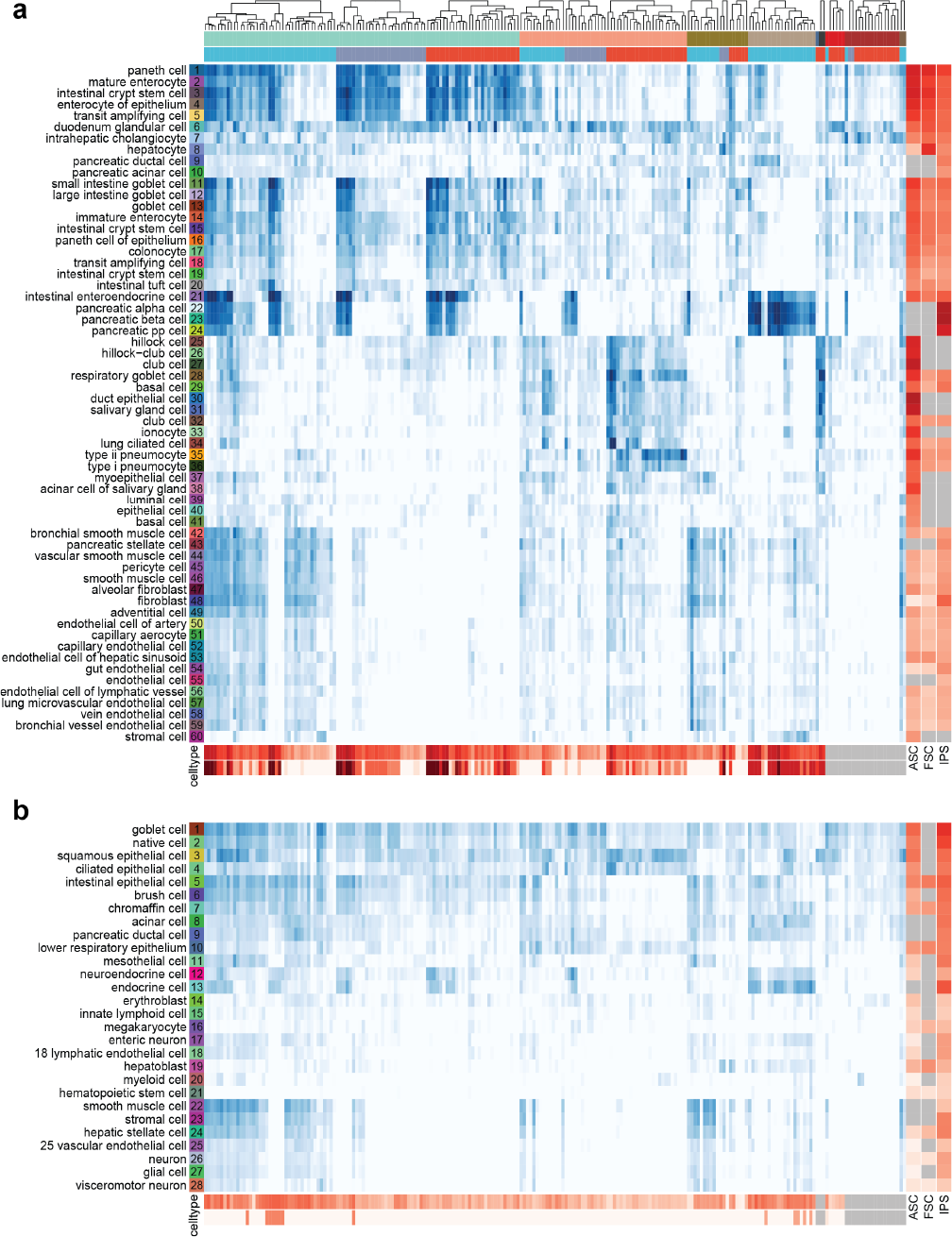
Organoid cell type comparison to cell types in fetal and adult primary tissue atlases. (a-b) Heatmaps display the similarity between each organoid sample and all adult primary tissue (a) and fetal primary tissue (b) cell types. The upper sidebar annotations indicate the organoid tissue and the source of stem cells. The right-hand sidebar annotations display the maximum similarity of different stem cell sources within their corresponding tissues. The lower sidebar annotations depict the average similarity of all cell types in corresponding tissues within each organoid sample.

**Figure S3.**
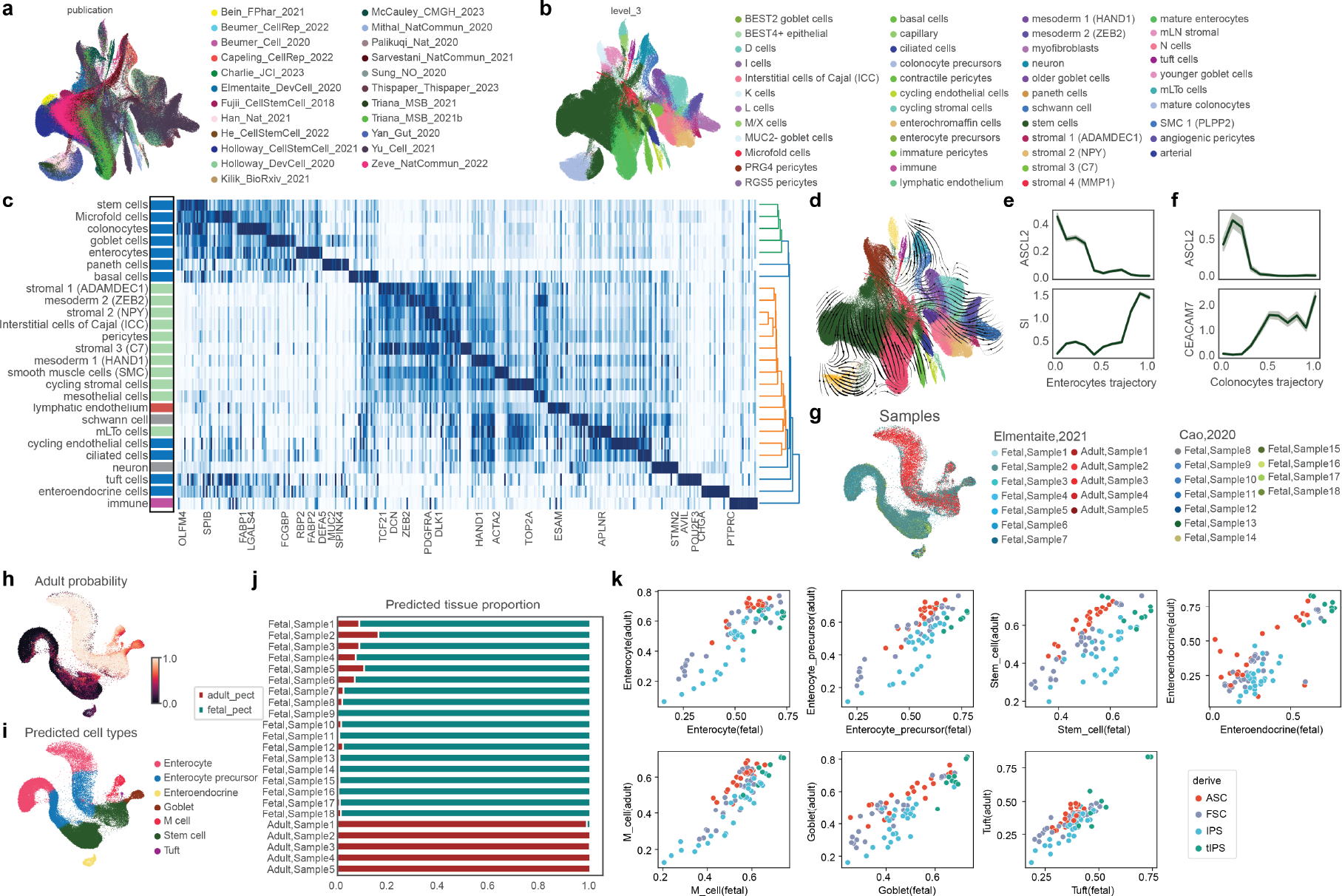
Integrated intestine organoid atlas and comparison to primary intestine tissue. (a-b) UMAP of the intestine organoid atlas colored by publications (a) and level 3 cell annotations (b). (c) Heatmap showing marker gene expression for each level 2 cell type in the intestine organoid atlas. Side stacked barplots show proportions of cell types at level 1 annotation. (d) UMAP of the integrated intestine organoid atlas with cells colored according to level 2 annotations. The stream arrows visualize the inferred velocity flow of cell states, providing insights into cellular dynamics. (e) Expression profiles along the pseudotime trajectory from stem cells to enterocytes of ASCL2 (stem cell) and SI (enterocytes). (f) Expression profiles along the pseudotime trajectory from stem cells to colonocytes of ASCL2 (stem cell) and CEACAM7 (colonocytes). (g-i) Projected 18 new fetal tissue samples and five new adult tissue samples, originating from two publications, onto fetal and adult primary tissue single-cell objects. The UMAP visualization displays cells colored according to different samples (g), adult probability (h), and predicted cell types (i). (j) Stacked barplots provide a visual representation of the predicted proportions of fetal or adult cells in all new tissue samples. (k) Scatter plots illustrate the maximum fetal or adult cell type similarity across all intestine organoid samples.

**Figure S4.**
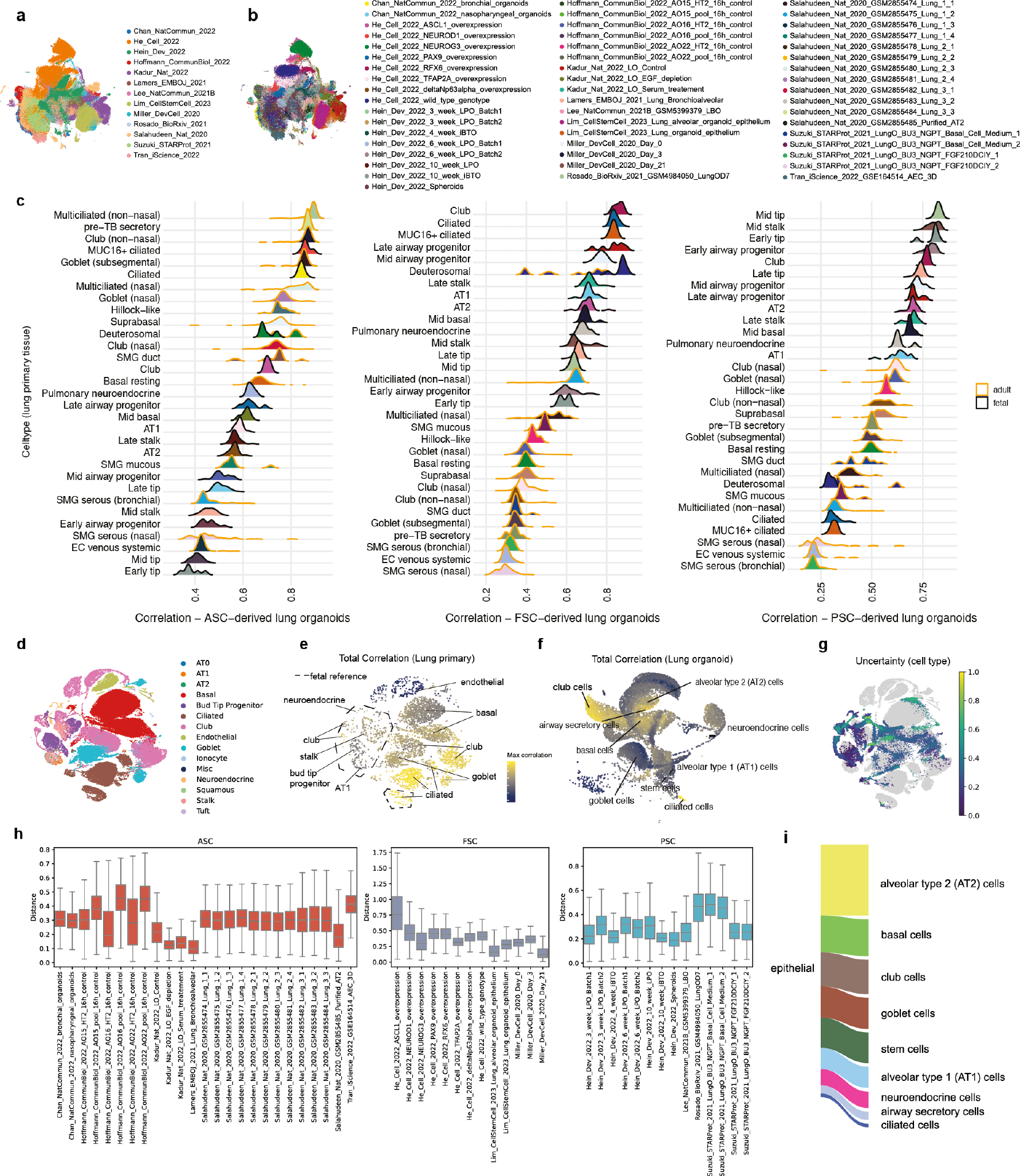
Lung organoid cell atlas integration, reference mapping and neighborhood correlation points to differences in PSC, FSC and ASC-organoid representation in primary tissue. (a) UMAP-representation of the HLOCA colored by included publications. (b) UMAP-representation of the HLOCA colored by included samples. (c) Ranked ridgeline plots per stem cell of origin, ordered by mean maximum correlation, showing max correlation distribution over neighborhoods between indicated primary cell type and ASC-, FSC- and PSC-organoids. (d) UMAP-representation of the integrated lung primary adult and fetal reference with simplified expert annotation based on original author annotation. (e) Neighborhood representation of the integrated lung primary adult and fetal reference embedding with coarse annotation from (d) showing maximum correlation to the HEOCA lung subset per neighborhood. (f) Neighborhood representation of the HEOCA lung subset embedding with level 2 annotation showing maximum correlation to the adult/fetal primary reference per neighborhood. (g) UMAP-representation colored by uncertainty of cell type prediction for lung organoid query cells mapped to the integrated lung primary adult and fetal reference. (h) Boxplot depicting distance of lung organoid query cells to the integrated lung primary adult and fetal reference by stem cell of origin and sample level resolution. (i) Composition of the HLOCA showing level 2 annotation ratio of epithelial cells.

**Figure S5.**
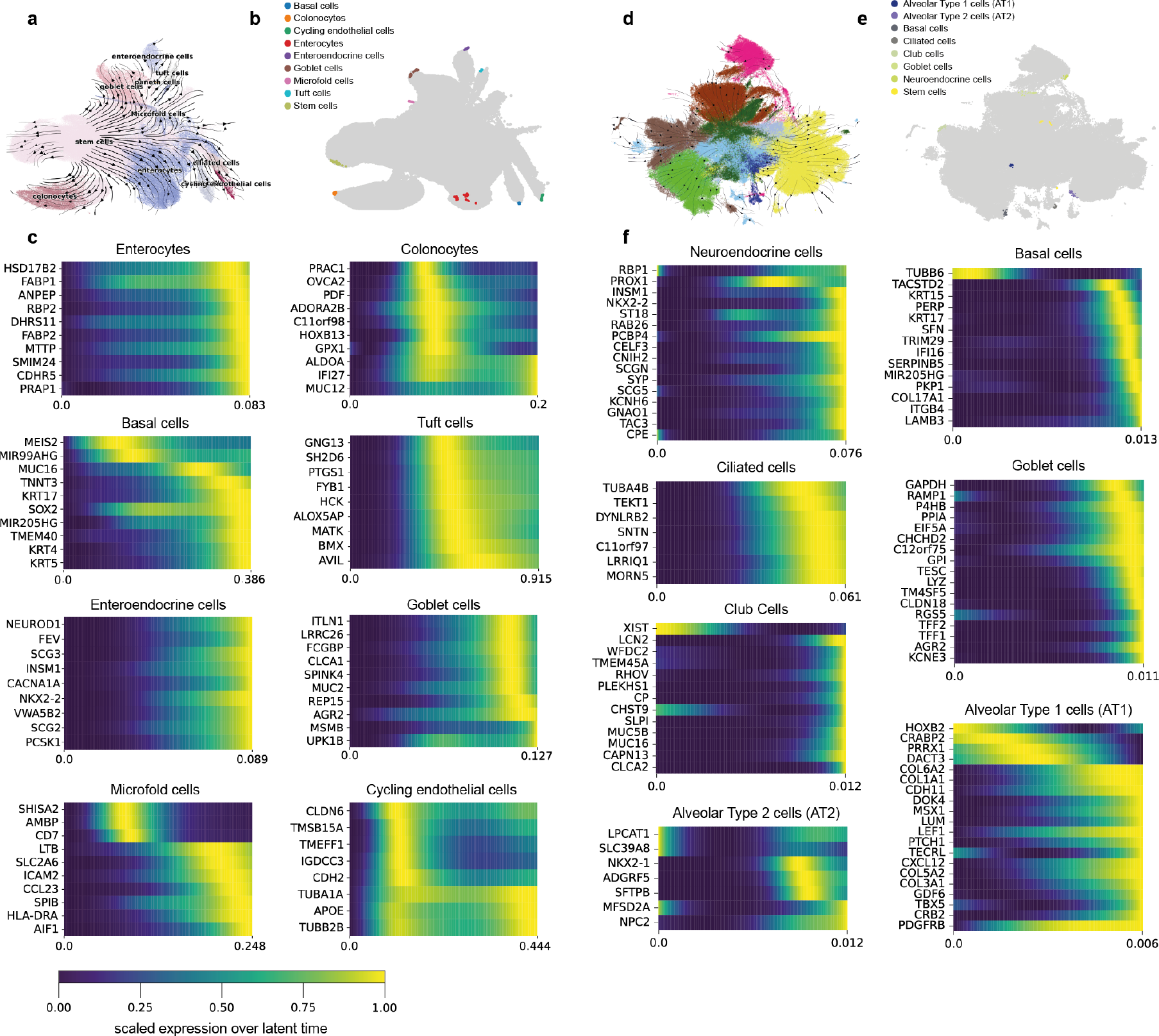
CellRank analysis of diffusion pseudotime reveals epithelial terminal states and trajectory-associated gene expression. (a) UMAP-representation of the HIOCA epithelial cells with diffusion pseudotime-dependent stream visualization of trajectories and level 2 cell type annotation. (b) UMAP-representation of the HIOCA epithelial cells with terminal states for each cell type pre-selected from macrostates generated through diffusion pseudotime by CellRank. (c) Cascade plots of top model fitting DPT trajectory associated gene expression peaks by terminal state per cell type (HIOCA, CellRank). (d) UMAP-representation of the HLOCA with diffusion pseudotime-dependent stream visualization of trajectories and level 2 cell type annotation. (e) UMAP-representation of the HLOCA with terminal states for each cell type pre-selected from macrostates generated through diffusion pseudotime by CellRank. (f) Cascade plots of top model fitting diffusion pseudotime trajectory associated gene expression peaks by terminal state per cell type (HLOCA, CellRank).

**Figure S6.**
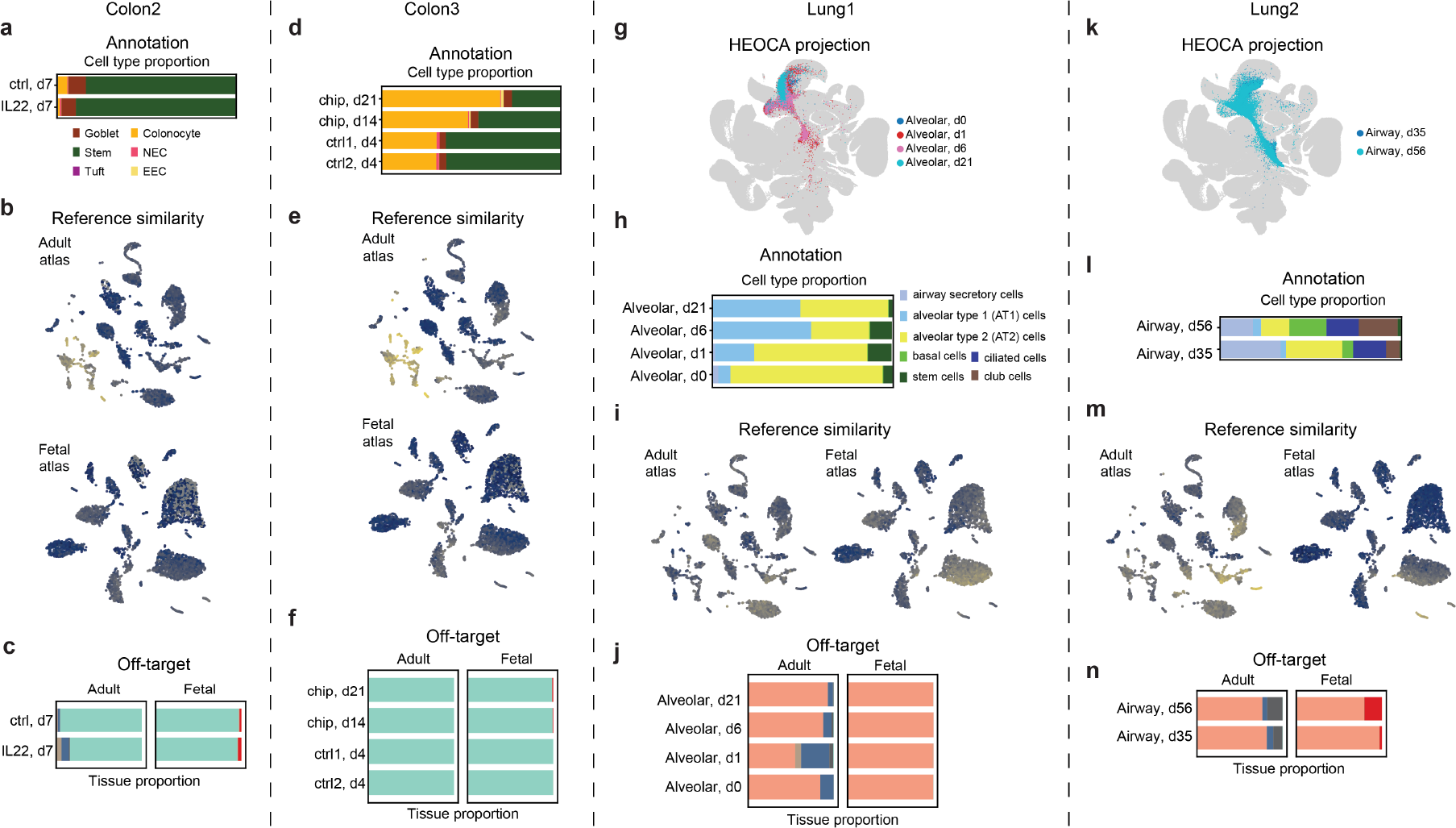
Assessment of new intestine and lung protocols through mapping to the organoid cell atlas. (a) Stacked barplot displays the predicted tissue proportions in new colon 2 protocol scRNA-seq samples. (b) UMAP representations of adult (top) and fetal (bottom) primary tissues are displayed, highlighting the maximum similarity achieved by new colon 2 protocol within the comprehensive cross-tissue atlas. (c) The barplots showing the tissue proportion of the most similar adult (left) and fetal (right) tissue across new colon 2 protocol time course samples. (d) Stacked barplot displays the predicted tissue proportions in new colon 3 protocol scRNA-seq samples. (e) UMAP representations of adult (top) and fetal (bottom) primary tissues are displayed, highlighting the maximum similarity achieved by new colon 3 protocol within the comprehensive cross-tissue atlas. (f) The barplots showing the tissue proportion of the most similar adult (left) and fetal (right) tissue across new colon 3 protocol time course samples. (g) UMAP of new lung 1 protocol time course scRNA-seq data mapped to the organoid atlas colored by different time points. (h) The barplot depicts the cell proportions of predicted level 2 cell types across new lung 1 protocol time course scRNA-seq data. (i) UMAP representations of adult (left) and fetal (right) primary tissues are displayed, highlighting the maximum similarity achieved by new lung 1 protocol within the comprehensive cross-tissue atlas. (j) The barplots showing the tissue proportion of the most similar adult (left) and fetal (right) tissue across new lung 1 protocol time course samples. (k) UMAP of new lung 2 protocol time course scRNA-seq data mapped to the organoid atlas colored by different time points. (l) The barplot depicts the cell proportions of predicted level 2 cell types across new lung 2 protocol time course scRNA-seq data. (m) UMAP representations of adult (left) and fetal (right) primary tissues are displayed, highlighting the maximum similarity achieved by new lung 2 protocol within the comprehensive cross-tissue atlas. (n) The barplots showing the tissue proportion of the most similar adult (left) and fetal (right) tissue across new lung 2 protocol time course samples.

**Figure S7.**
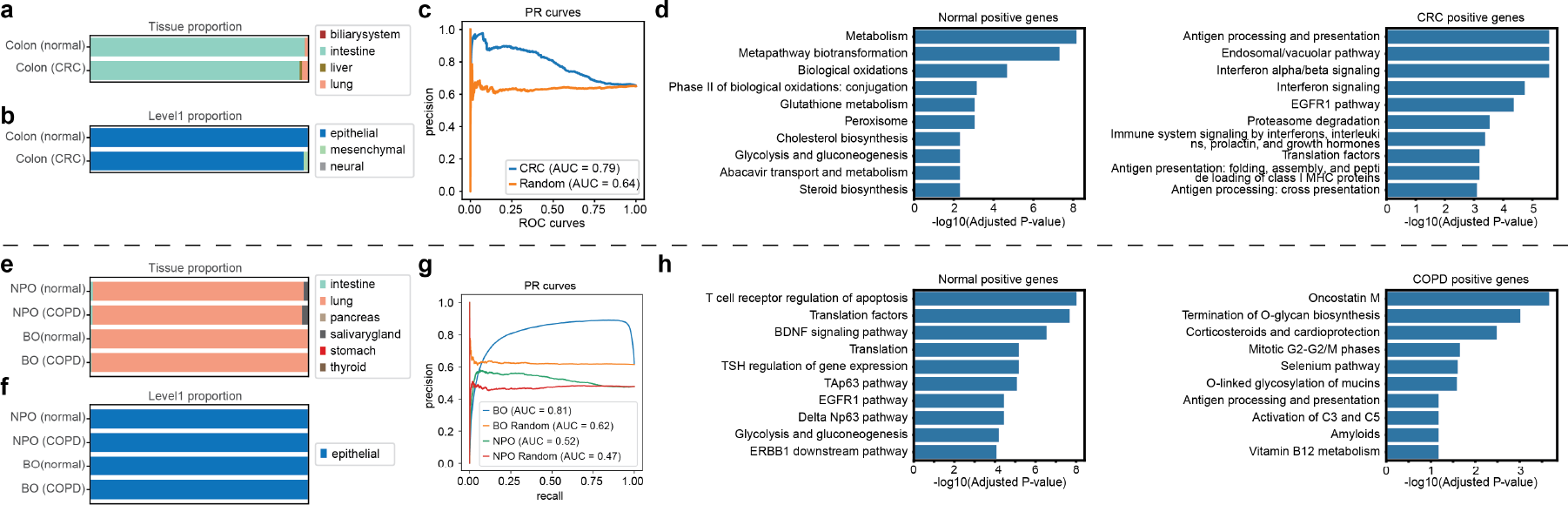
Mapping organoid disease model data to the HEOCA reveals differences in distance to atlas cell types and alterations in expression programs. (a) Stacked barplot represents the predicted tissue proportions within scRNA-seq samples from both normal and cancerous colon tissues. (b) A stacked barplot is employed to illustrate the predicted proportions of level 2 cell types in scRNA-seq samples obtained from normal and cancerous colon tissues. (c) A PR plot shows the predicted colorectal cancer cells based on the mean distance between single cells and their nearest neighbors within the HEOCA atlas. (d) The barplots provided in this section showcase pathways that exhibit significant enrichment in colonocytes for both normal and cancerous colon tissue samples. (e) A stacked barplot displays the predicted tissue proportions within scRNA-seq samples from both normal and COPD lung samples. (f) The stacked barplot illustrates the predicted proportions of level 2 cell types in scRNA-seq samples obtained from normal and COPD lung organoid samples. (g) The PR plot shows the predicting COPD lung cells, relying on the mean distance between single cells and their nearest neighbors within the HEOCA atlas. (h) Barplots depict pathways significantly enriched in basal cells for both normal and BO COPD lung organoid samples.

## Notes

### Competing Interest Statement

F.J.T. consults for Immunai Inc., Singularity Bio B.V., CytoReason Ltd, Cellarity, and has ownership interest in Dermagnostix GmbH and Cellarity. Other authors declare no conflict of interest.

### Summary of Updates

No

